# Inter- and intra-animal variation of integrative properties of stellate cells in the medial entorhinal cortex

**DOI:** 10.1101/678565

**Authors:** Hugh Pastoll, Derek Garden, Ioannis Papastathopoulos, Gülşen Sürmeli, Matthew F. Nolan

## Abstract

Distinctions between cell types underpin organisational principles for nervous system function. Functional variation also exists between neurons of the same type. This is exemplified by correspondence between grid cell spatial scales and synaptic integrative properties of stellate cells (SCs) in the medial entorhinal cortex. However, we know little about how functional variability is structured either within or between individuals. Using ex-vivo patch-clamp recordings from up to 55 SCs per mouse, we find that integrative properties vary between mice and, in contrast to modularity of grid cell spatial scales, have a continuous dorsoventral organisation. Our results constrain mechanisms for modular grid firing and provide evidence for inter-animal phenotypic variability among neurons of the same type. We suggest that neuron type properties are tuned to circuit level set points that vary within and between animals.

## Introduction

The concept of cell types provides a general organising principle for understanding biological structures including the brain (Regev et al., 2017; Zeng and Sanes, 2017). The simplest conceptualisation of a neuronal cell type, as a population of phenotypically similar neurons with features that cluster around a single set point (Wang et al., 2011b), is extended by observations of variability in cell type features, suggesting that some neuronal cell types may be conceived as clustering along a line rather than a point in a feature space (Cembrowski and Menon, 2018; O’Donnell and Nolan, 2011)(Figure 1A). Correlations between the functional organisation of sensory, motor and cognitive circuits and electrophysiological properties of individual neuronal cell types suggest that this feature variability underlies key neural computations (Adamson et al., 2002; Angelo et al., 2012; Fletcher and Williams, 2018; Garden et al., 2008; Giocomo et al., 2007; Kuba et al., 2005; O’Donnell and Nolan, 2011). However, within cell type variability has typically been deduced by combining data obtained from multiple animals. In contrast, the structure of variation within individual animals or between different animals has received little attention. For example, apparent clustering of properties along lines in feature space could reflect a continuum of set points, or could result from a small number of discrete set points that are obscured by inter-animal variation (Figure 1B). Moreover, while investigations of invertebrate nervous systems show that set points may differ between animals (Goaillard et al., 2009), it is not clear whether mammalian neurons exhibit similar phenotypic diversity (Figure 1B). Distinguishing these possibilities requires many more electrophysiological observations per animal than are obtained in typical studies. Although methods for high quality recording from SCs in ex-vivo brain slices are well established (Pastoll et al., 2012b), in previous studies typically fewer than 5 recordings per animal have been made, which is much less than our estimate of the minimum number of observations required to test for modularity.

**Figure 1.**
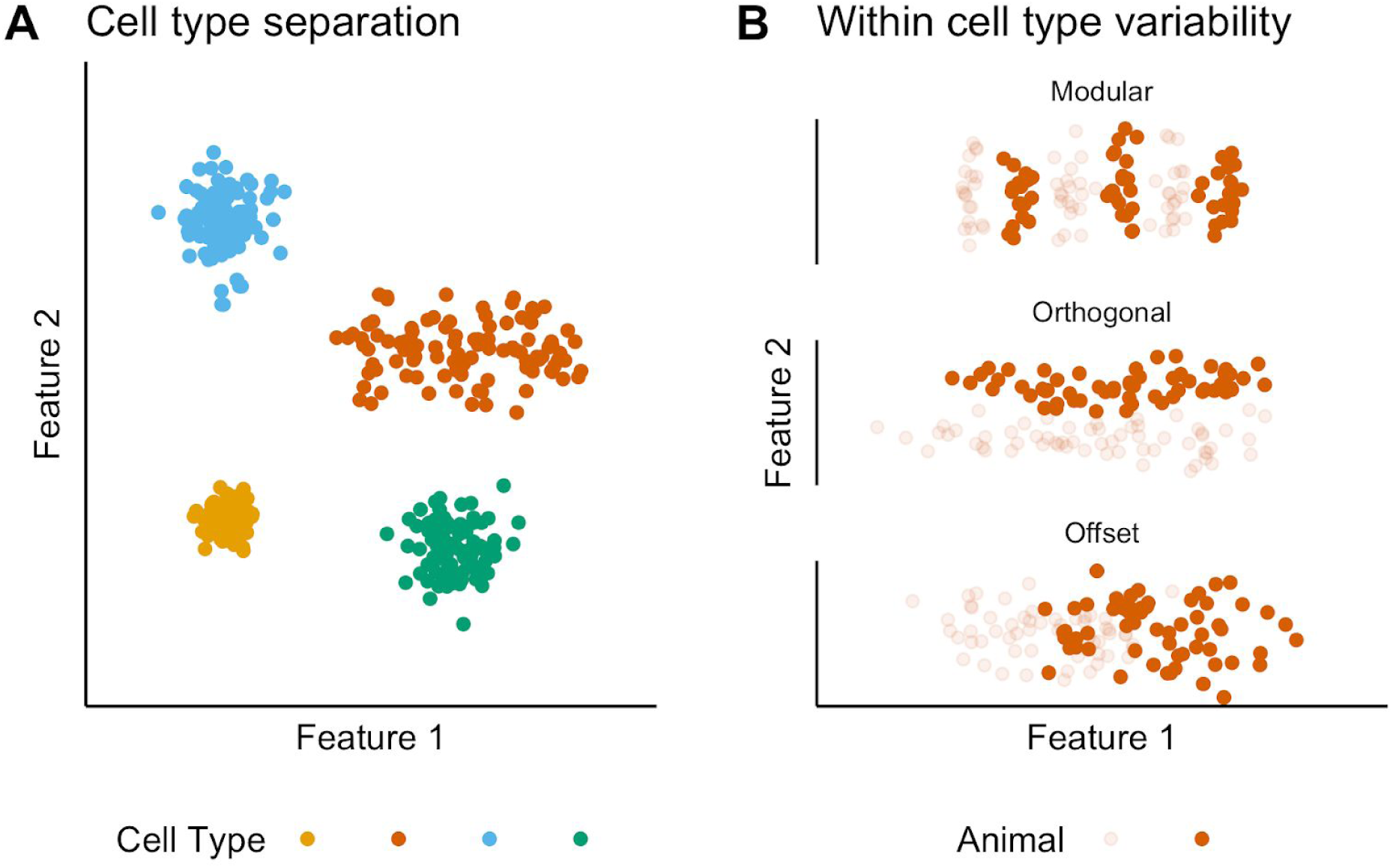
Classification and variability of neuronal cell types. (A) Neuronal cell types are identifiable by features clustering around a distinct point (blue, green and yellow) or a line (red) in feature space. The clustering implies that neuron types are defined by either a single set point (blue, green and yellow) or by multiple set points spread along a line (red). (B) Phenotypic variability of a single neuron type could result from distinct set points that subdivide the neuron type and appear continuous when data from multiple animals are combined (modular), from differences in components of a set point that do not extend along a continuum but that in different animals cluster at different locations in feature space (orthogonal), or from differences between animals in the range covered by a continuum of set points (offset). These distinct forms of variability can only be made apparent by measuring features from many neurons from multiple animals (colours).

Stellate cells in layer 2 (SCs) of the medial entorhinal cortex (MEC) provide a striking example of correspondence between functional organisation of neural circuits and variability of electrophysiological features within a single cell type. The MEC contains neurons that encode an animal’s location through grid-like firing fields (Fyhn et al., 2004). The spatial scale of grid fields follows a dorsoventral organisation (Hafting et al., 2005), which is mirrored by a dorsoventral organisation in key electrophysiological features of SCs (Boehlen et al., 2010; Dodson et al., 2011; Garden et al., 2008; Giocomo and Hasselmo, 2008a; Giocomo et al., 2007; Pastoll et al., 2012a). Grid cells are organised into discrete modules (Stensola et al., 2012), cells in a module have a similar grid scale and orientation (Barry et al., 2007; Gu et al., 2018; Stensola et al., 2012) and progressively more ventral modules are composed of cells with wider grid spacing (Stensola et al., 2012). Studies that demonstrate dorsoventral organisation of integrative properties of SCs have so far relied on pooling of relatively few measurements per animal. Hence, it is unclear whether organisation of these cellular properties is modular, as one might expect if they directly set the scale of grid firing fields in individual grid cells (Giocomo et al., 2007). The possibility that set points for electrophysiological properties of SCs differ between animals has also not previously been considered.

Evaluation of variability between and within animals requires statistical approaches not typically used in single-cell electrophysiological investigations. Given appropriate assumptions, inter-animal differences can be assessed using mixed effect models that are well established in other fields (Baayen et al., 2008; Geiler-Samerotte et al., 2013). Because tests of whether data arise from modular as opposed to continuous distributions have received less general attention, to facilitate detection of modularity using relatively few observations we introduce a modification of the gap statistic algorithm (Tibshirani et al., 2001) that estimates the number of modes in a dataset while controlling for observations expected by chance (see Methods and Supplemental Figures 1 - 5). This algorithm performs well compared with other measures (Giocomo et al., 2014; Stensola et al., 2012), which we find are prone to high false positive rates (Supplemental Figure 4A). We find that recordings from approximately 30 SCs per animal should be sufficient to detect modularity using the modified gap statistic algorithm and given the experimentally observed separation between grid modules (see Methods and Supplemental Figures 2-3).

**Figure 2.**
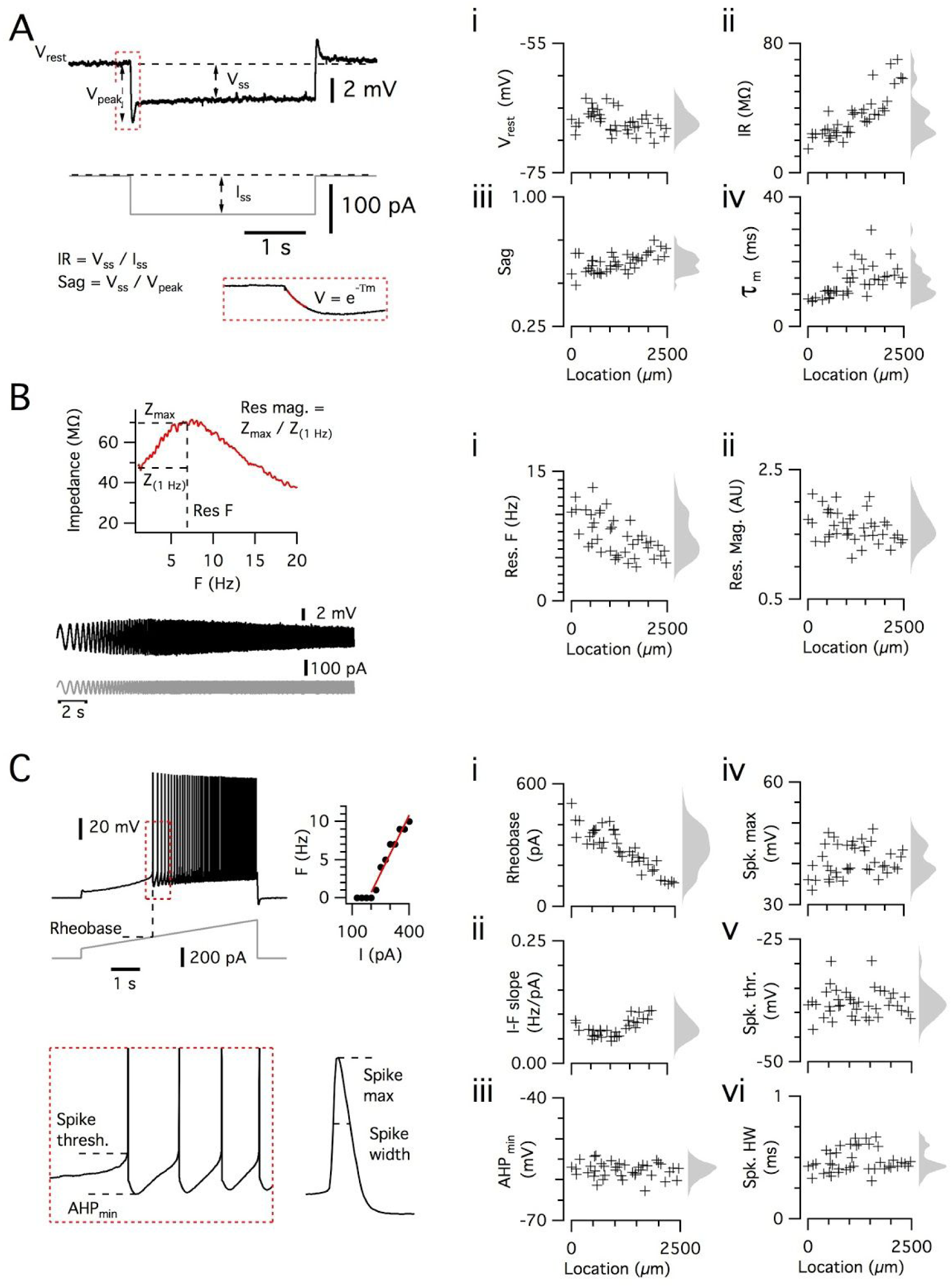
Dorsoventral organization of intrinsic properties of stellate cells from a single animal. (A-C) Waveforms (grey traces) and example responses (black traces) from a single mouse for step (A), ZAP (B) and ramp (C) stimuli (left). Properties derived from each protocol are shown plotted against recording location (each data point is a black cross)(right). KSDs with arbitrary bandwidth are displayed to the right of each data plot to facilitate visualisation of the property’s distribution when location information is disregarded (light grey pdfs). (A) Injection of a series of current steps enables measurement of the resting membrane potential (Vrest) (i), the input resistance (IR) (ii), the sag coefficient (sag) (iii) and the membrane time constant (Τm) (iv). (B) Injection of ZAP current waveform enables calculation of an impedance amplitude profile, which was used to estimate the resonance resonant frequency (Res. F) (i) and magnitude (Res. mag) (ii). (C) Injection of a slow current ramp enabled measurement of the rheobase (i), the slope of the current-frequency relationship (I-F slope) (ii), and using only the first 5 spikes in each response (enlarged zoom, lower left) the AHP minimum value (AHP_min_) (iii), the spike maximum (Spk. max) (iv), the spike width at half height (Spk. HW) (v) and the spike threshold (Spk. thr.) (vi).

**Figure 3.**
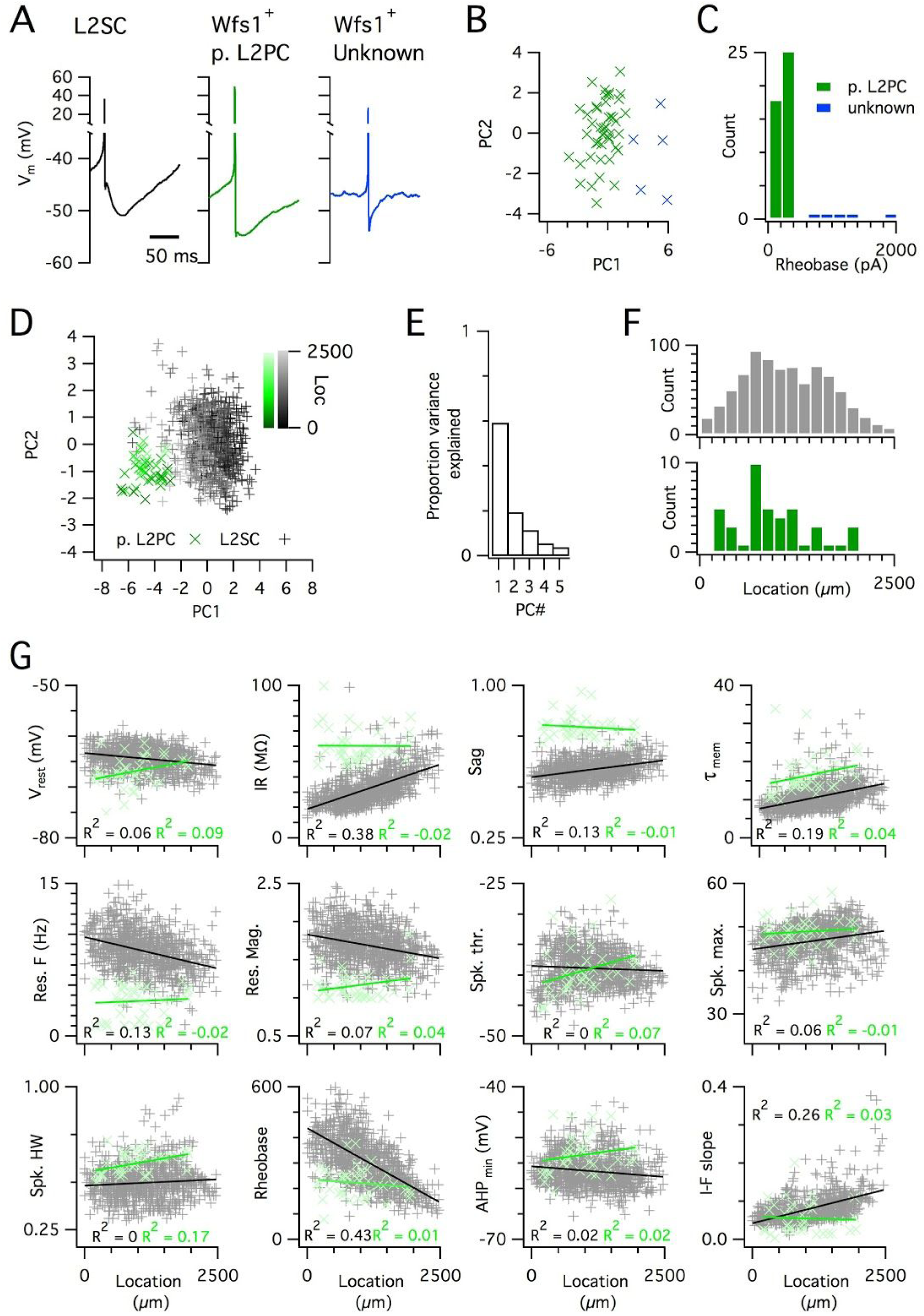
Distinct and dorsoventrally organised properties of layer 2 stellate cells. (A) Representative action potential afterhyperpolarization waveforms from a SC (left), a pyramidal cell (middle) and an unidentified cell (right). The pyramidal and unidentified cells were both positively labelled in Wfs1^Cre^ mice. (B) Plot of the first versus the second principal component from PCA of the properties of labelled neurons in Wfs1^Cre^ mice reveals two populations of neurons. (C) Histogram showing the distribution of rheobase values of cells positively labelled in Wfs1^Cre^ mice. The two groups identified in B can be distinguished by their rheobase. (D) Plot of the first two principal components from PCA of the properties of the L2PC (n = 44, green) and SC populations (n = 840, black). Putative pyramidal cells (x) and SCs (+) are colored according to their dorsoventral location (inset shows the scale). (E) Proportion of total variance explained by the first five principal components for the analysis in (D). (F) Histograms of locations of recorded SCs (upper) and L2PCs (lower). (G) All values of measured features from all mice are plotted as a function of the dorsoventral location of the recorded cells. Lines indicate fits of a linear model to the complete datasets for SCs (black) and L2PCs (green). Putative pyramidal cells (x, green) and SCs (+, black). Adjusted R^2^ values use the same colour scheme.

We set out to establish the nature of the set points that establish integrative properties of SCs by measuring intra- and inter-animal variation in key electrophysiological features using experiments that maximise the number of SCs recorded per animal. Our results suggest that set points for individual features of a neuronal cell type are established at a population level, differ between animals and follow a continuous organisation.

## Results

### Sampling integrative properties from many neurons per animal

Before addressing intra- and inter-animal variability we first describe the data set used for the analyses that follow. We established procedures to facilitate recording of integrative properties of many SCs from a single animal (see Methods). With these procedures, we measured and analysed electrophysiological features of 836 SCs (n/mouse: range 11-55; median = 35) from 27 mice (median age = 37.4 days, age range = 18 - 57 days). The mice were housed either in a standard home cage (N = 18 mice, n = 583 neurons) or from postnatal day 16 in a 2.4 × 1.2 m cage, which provided a large environment that could be freely explored (N = 9, n = 253, median age = 38 days)(Supplemental Figure 6). For each neuron we measured six sub-threshold integrative properties (Figure 2A-B) and six supra-threshold integrative properties (Figure 2C). Until indicated otherwise we report analysis of datasets that combine the groups of mice housed in standard and large home cages and that span the full range of ages.

Because SCs are found intermingled with pyramidal cells in layer 2 (L2PCs), and as misclassification of L2PCs as SCs would likely confound investigation of intra-SC variation, we validated our criteria for distinguishing each cell type. To establish characteristic electrophysiological properties of L2PCs we recorded from neurons in layer 2 identified by Cre-dependent marker expression in a Wfs1^Cre^ mouse line (Sürmeli et al., 2015). Expression of Cre in this line, and a similar line (Kitamura et al., 2014), labels pyramidal cells in layer 2 (L2PCs) that project to the CA1 region of the hippocampus, but does not label SCs (Kitamura et al., 2014; Sürmeli et al., 2015). We identified two populations of neurons in layer 2 of MEC that were labelled in Wfs1^Cre^ mice (Figure 3A-C). The more numerous population had properties consistent with L2PCs (Figure 3A, G) and could be separated from the unidentified population on the basis of a lower rheobase (Figure 3C). The unidentified population had firing properties typical of layer 2 interneurons (cf. (Gonzalez-Sulser et al., 2014)). A principal component analysis (PCA)(Figure 3D-F) clearly separated the L2PC population from the SC population, but did not identify subpopulations of SCs. The less numerous population has properties similar to those of inhibitory interneurons and also clearly distinct from SCs (Figure 3A, C). These data demonstrate that the SC population used for our analyses is distinct from excitatory pyramidal cells also found in layer 2 of the MEC.

To further validate the large SC dataset we assessed the location-dependence of individual features, several of which have previously been found to depend on the dorso-ventral location of the recorded neuron (Boehlen et al., 2010; Booth et al., 2016; Garden et al., 2008; Giocomo et al., 2007; Pastoll et al., 2012a). We initially fit the dependence of each feature on dorsoventral position using a standard linear regression model. We found substantial (adjusted R^2^ > 0.1) dorsoventral gradients in input resistance, sag, membrane time constant, resonant frequency, rheobase and the current-frequency (I-F) relationship (Figure 3G). In contrast to SCs, we did not find evidence for dorsoventral organisation of these features in L2PCs (Figure 3G). Thus, our large dataset replicates the previously observed dependence of integrative properties of SCs on their dorsoventral position, and shows that this location dependence further distinguishes SCs from L2PCs.

### Inter-animal differences in intrinsic properties of stellate cells

To what extent does variability between the integrative properties of SCs at a given dorsoventral location (Figure 3G) arise from differences between animals? Comparing specific features from individual animals suggested that their distributions could be almost completely non-overlapping despite consistent and strong dorsoventral tuning (Figure 4A). If this apparent inter-animal variability results from random sampling of a distribution determined by a common underlying set point, then fitting the complete data set with a mixed model in which animal identity is included as a random effect should reconcile the apparent differences between animals (Figure 4B). In this scenario, the conditional R^2^ estimated from the mixed model, in other words the estimate of variance explained by animal identity and location, should be similar to the marginal R^2^ value, which indicates the variance explained by location only. In contrast, if differences between animals contribute to experimental variability, the mixed model should predict different fitting parameters for each animal, and the estimated conditional R^2^ should be greater than the corresponding marginal R^2^ (Figure 4C).

**Figure 4.**
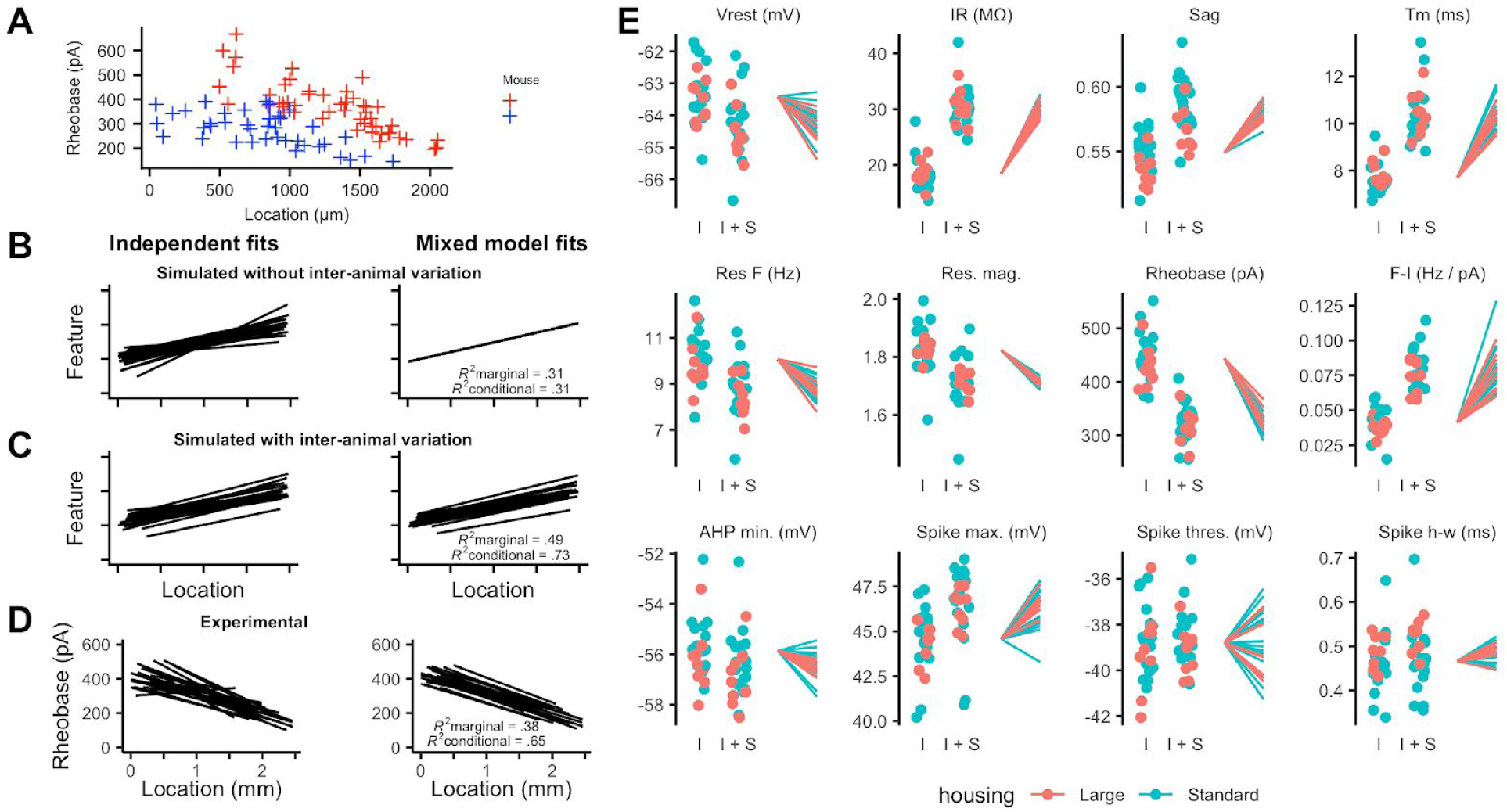
Inter-animal variability and dependence on environment of intrinsic properties of stellate cells. (A) Examples of rheobase as a function of dorsoventral position for two mice. (B-C) Each line is the fit of simulated data from a different subject for datasets in which there is no inter-subject variability (B) or in which intersubject variability is present (C). Fitting data from each subject independently with linear regression models suggests intersubject variation in both datasets (left). In contrast, after fitting mixed effect models (right) intersubject variation is no longer suggested for dataset in which it is absent (B) but remains for the dataset in which it is present (C). (D) Each line is the fit of rheobase as a function of dorsoventral location for a single mouse. The fits were carried out independently for each mouse (left) or using a mixed effect model with mouse identity as a random effect (right). (E) The intercept (I), sum of the intercept and slope (I + S), and slopes realigned to a common intercept (right) for each mouse obtained by fitting mixed effect models for each property as a function of dorsoventral position.

Fitting the experimental measures for each feature with mixed models suggests that differences between animals contribute substantially to the variability in properties of SCs. In contrast to simulated data in which inter-animal differences are absent (Figure 4B), differences in fits between animals remained after fitting with the mixed model (Figure 4D). This corresponds with expectations from fits to simulated data containing inter-animal variability (cf. Figures 4C). To visualise inter-animal variability for all measured features we plot for each animal the intercept of the model fit (I), the predicted value at a location 1 mm ventral from the intercept (I+S) and the slope (lines)(Figure 4E). Strikingly, even for features such as rheobase and input resistance (IR) that are highly tuned to a neurons’ dorsoventral position, the extent of variability between animals is similar to the degree to which the property changes between dorsal and mid-levels of the MEC.

If set points that determine integrative properties of SCs do indeed differ between animals, then mixed models should provide a better account of the data than linear models generated by pooling data across all animals. Consistent with this, we found that mixed models for all electrophysiological features gave a substantially better fit to the data than linear models that considered all neurons as independent (adjusted p < 2 × 10^-17^ for all models, *χ*^2^ test, Table 1). Furthermore, even for properties with substantial (R^2^ value > 0.1) dorsoventral tuning, the conditional R^2^ value for the mixed effect model was substantially larger than the marginal R^2^ value (Figure 4D and Table 1). Together, these analyses demonstrate inter-animal variability in key electrophysiological features of SCs suggesting that set points that establish the underlying integrative properties differ between animals.

**Table 1.**
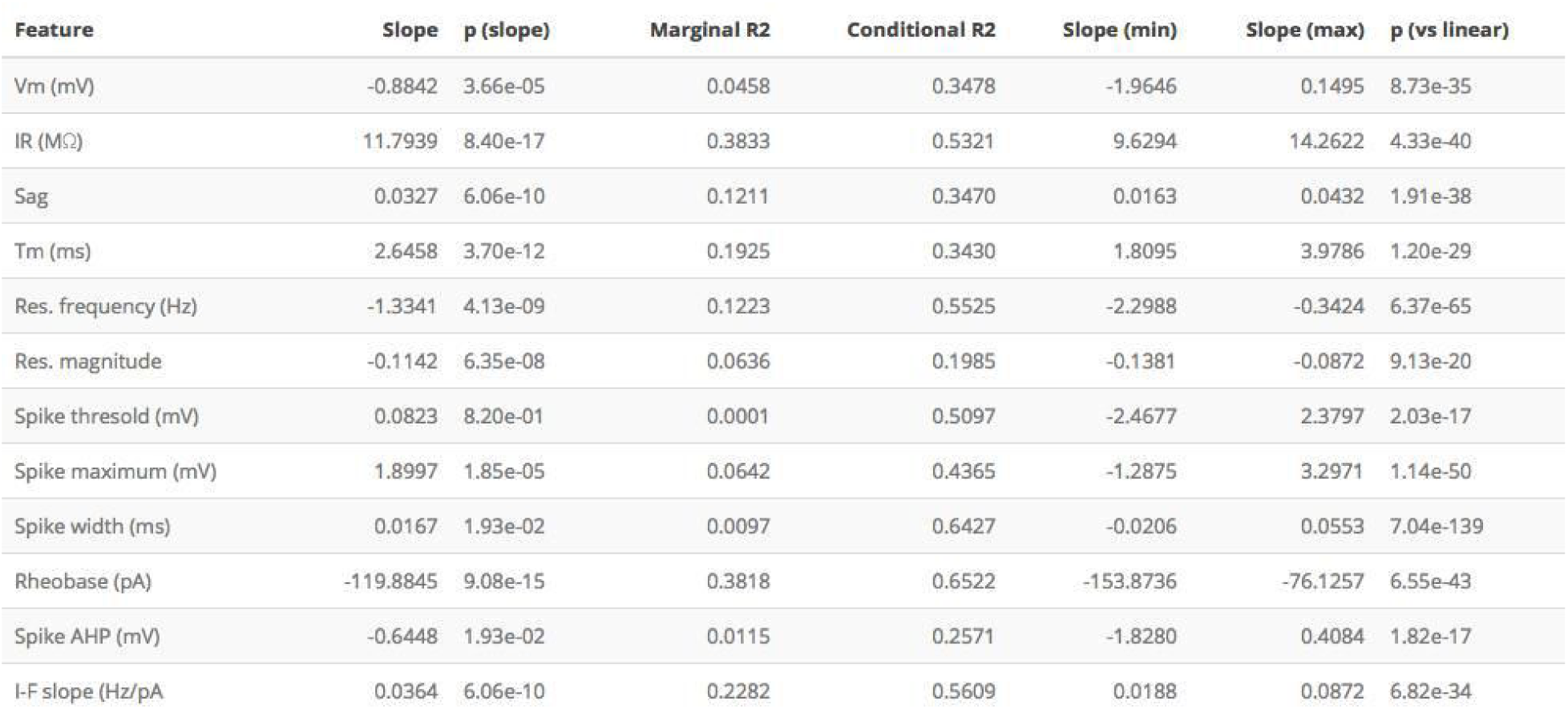
Dependence of electrophysiological features of SCs on dorsoventral position. Key statistical parameters from analyses of the relationship between each measured electrophysiological feature and dorsoventral location. Slope is the population slope from fitting a mixed effect model for each feature with location as a fixed effect, with p(slope) obtained by comparing this model to a model without location as a fixed effect (*χ*^2^ test). The marginal and conditional R^2^ values, and the minimum and maximum slopes across all mice, are obtained from the fits of mixed effect models that contain location as a fixed effect. The estimate p(vs linear) is obtained by comparing the mixed effect model containing location as a fixed effect with a corresponding linear model without random effects (*χ*^2^ test). The values of p(slope) and p(vs linear) were adjusted for multiple comparisons using the method of Benjamini and Hochberg (Benjamini and Hochberg, 1995). Units for the slope measurements are units for the property mm^-1^. The data are from 27 mice.

### Experience-dependence of intrinsic properties of stellate cells

Because neuronal integrative properties may be modified by changes in neural activity (Zhang and Linden, 2003), we asked if experience influences the measured electrophysiological features of SCs. We reasoned that modifying the space through which animals can navigate may drive experience-dependent plasticity in the MEC. As standard mouse housing has dimensions less than the distance between firing fields of more ventrally located grid cells (Brun et al., 2008; Hafting et al., 2005), we tested whether electrophysiological features of SCs differ between mice housed in larger environments (28800 cm^2^) compared with standard home cages (530 cm^2^).

We compared the mixed models described above to models in which housing was also included as a fixed effect. To minimise the effects of age on SCs (Boehlen et al., 2010; Burton et al., 2008)(Supplemental Table 2), we focussed these and subsequent analyses on mice between P33 and P44 (N = 25, n = 779). We found that larger housing was associated with a smaller sag coefficient indicating an increased sag response, a lower resonant frequency and a larger spike half-width (adjusted p < 0.05; Figure 4E, Supplemental Table 3, Supplemental analyses). These differences were primarily from changes to the magnitude rather than the location-dependence of each feature. Other electrophysiological features appeared unaffected by housing.

To determine whether inter-animal differences remain after accounting for housing we compared mixed models that include dorsoventral location and housing as fixed effects with equivalent linear regression models in which individual animals are not accounted for. Mixed models incorporating animal identity continued to provide a better account of the data, both for features that were dependent on housing (adjusted p < 2.8 × 10^-21^) and for features that were not (adjusted p < 1.4 × 10^-7^)(Supplemental Table 4).

Together, these data suggest that specific electrophysiological features of SCs may be modified by experience of large environments. After accounting for housing, significant inter-animal variation remains, suggesting that additional mechanisms acting at the level of animals rather than individual neurons also determine differences between SCs.

### Inter-animal differences remain after accounting for additional experimental parameters

To address the possibility that other experimental or biological variables could contribute to inter-animal differences, we evaluated the effects of home cage size (Supplemental Tables 3-4), brain hemisphere (Supplemental Table 5), mediolateral position (Supplemental Table 6), the identity of the experimenter (Supplemental Table 7) and time since slice preparation (Supplemental Tables 8 and 9). Several of the variables influenced some measured electrophysiological features, for example properties primarily related to the action potential waveform depended on mediolateral position of the recorded neuron (Supplemental Table 6)(cf. (Canto and Witter, 2012)), but significant inter-animal differences remained after accounting for each variable. We carried out further analyses using models that included housing, mediolateral position, experimenter identity and the direction in which sequential recordings were obtained as fixed effects (Supplemental Table 10), using models fit to minimal datasets in which housing, mediolateral position and the recording direction were identical (Supplemental Table 11), and to consider the possible impact of tissue variability (Supplemental Tables 12 and 13). These analyses again found evidence for significant inter-animal differences. We also note that conditional R^2^ values of electrophysiological features of L2PCs are much lower than for SCs (cf. Table 1 and Supplemental Table 1) suggesting that inter-animal variation may be specific to SCs, and further arguing that inter-animal differences result from biological rather than technical variability. Together, these analyses indicate that differences between animals remain after accounting for experimental and technical factors that might contribute to variation in measured features of SCs.

### The distribution of intrinsic properties is consistent with a continuous rather than a modular organisation

The dorsoventral organisation of SC integrative properties is well established, but whether this results from within animal variation consistent with a small number of discrete set points that underlie a modular organisation (cf. Figure 1B) is unclear. To evaluate modularity we used datasets with n ≥ 34 (N = 15, median age = 37 days, age range = 18 - 43 days). We focus initially on rheobase, which is the property with strongest correlation to dorsoventral location, and resonant frequency, which is related to the oscillatory dynamics underlying dorsoventral tuning in some models of grid firing. For n ≥ 34 we expect that if properties are modular then this would be detected by the modified gap statistic in at least 50% of animals (Supplemental Figures 2C, 3). In contrast, we find that for datasets from the majority of animals the modified gap statistic identifies only a single mode in the distribution of rheobase values (Figure 5A and Figure 6)(N = 13/15) and of resonant frequencies (Figure 5B and Figure 6) (N = 14/15), indicating that these properties have a continuous rather than a modular distribution. Consistent with this, smoothed distributions did not show clearly separated peaks for either property (Figure 5). The mean and 95% confidence interval for the probability of evaluating a dataset as clustered (p_detect_), was 0.133 and 0.02–0.4 for rheobase and 0.067 and 0.002–0.32 for resonant frequency. These values of p_detect_ were not significantly different from the proportions expected from the complete absence of clustering (p = 0.28 and 0.66, binomial test). Thus, the rheobase and resonant frequency of SCs, while depending strongly on a neuron’s dorsoventral position, do not have a detectable modular organisation.

**Figure 5.**
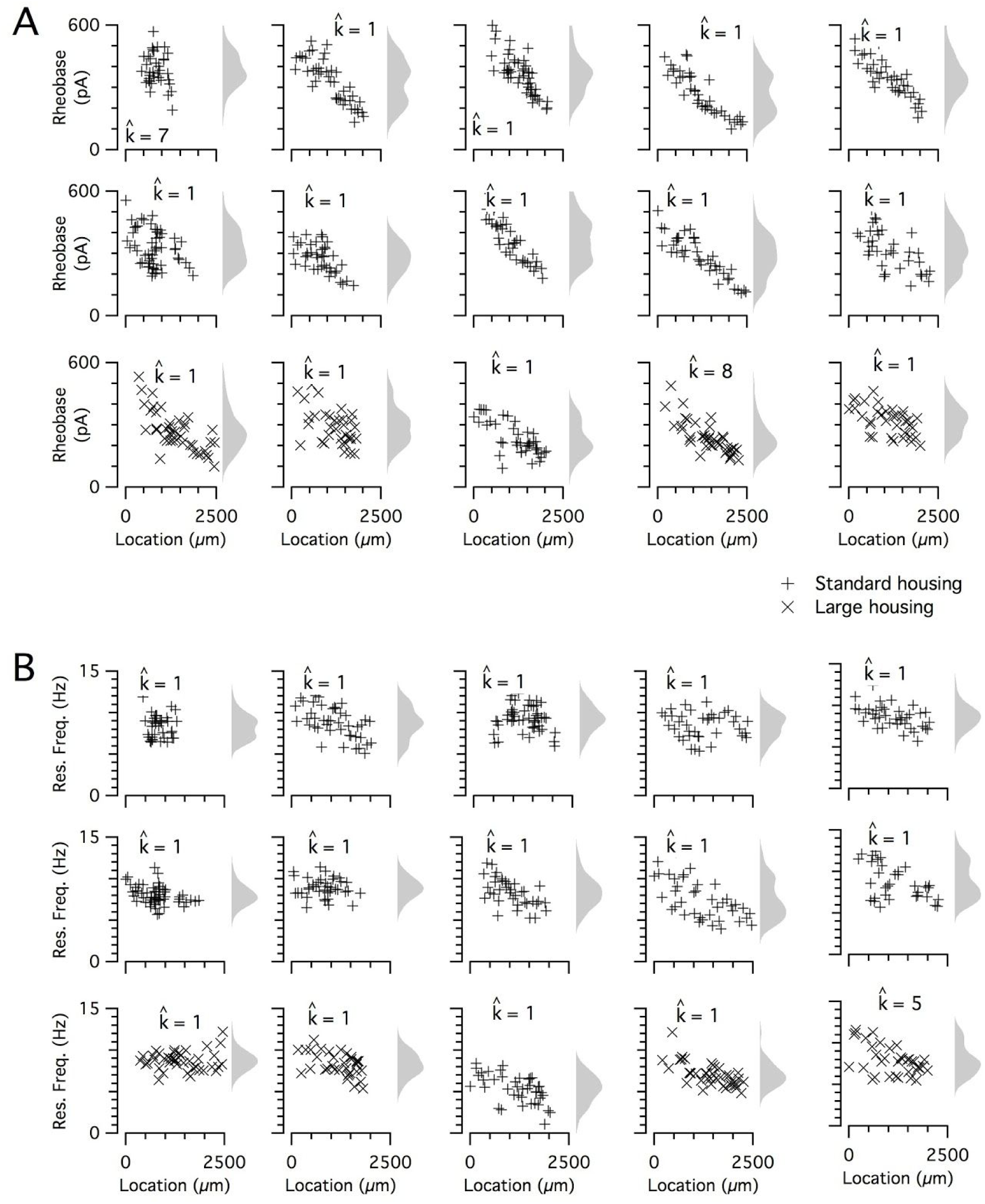
Input resistance and resonant frequency do not have a detectable modular organisation. (A-B) Input resistance (A) and resonant frequency (B) are plotted as a function of dorsoventral position separately for each every animal. Marker colour and types indicate the age and housing conditions of the animal (‘+’s standard housing, ‘x’s large housing). KSDs (arbitrary bandwidth, same for all animals) are also shown. The number of clusters in the data (k_est_) is indicated for each animal.

**Figure 6.**
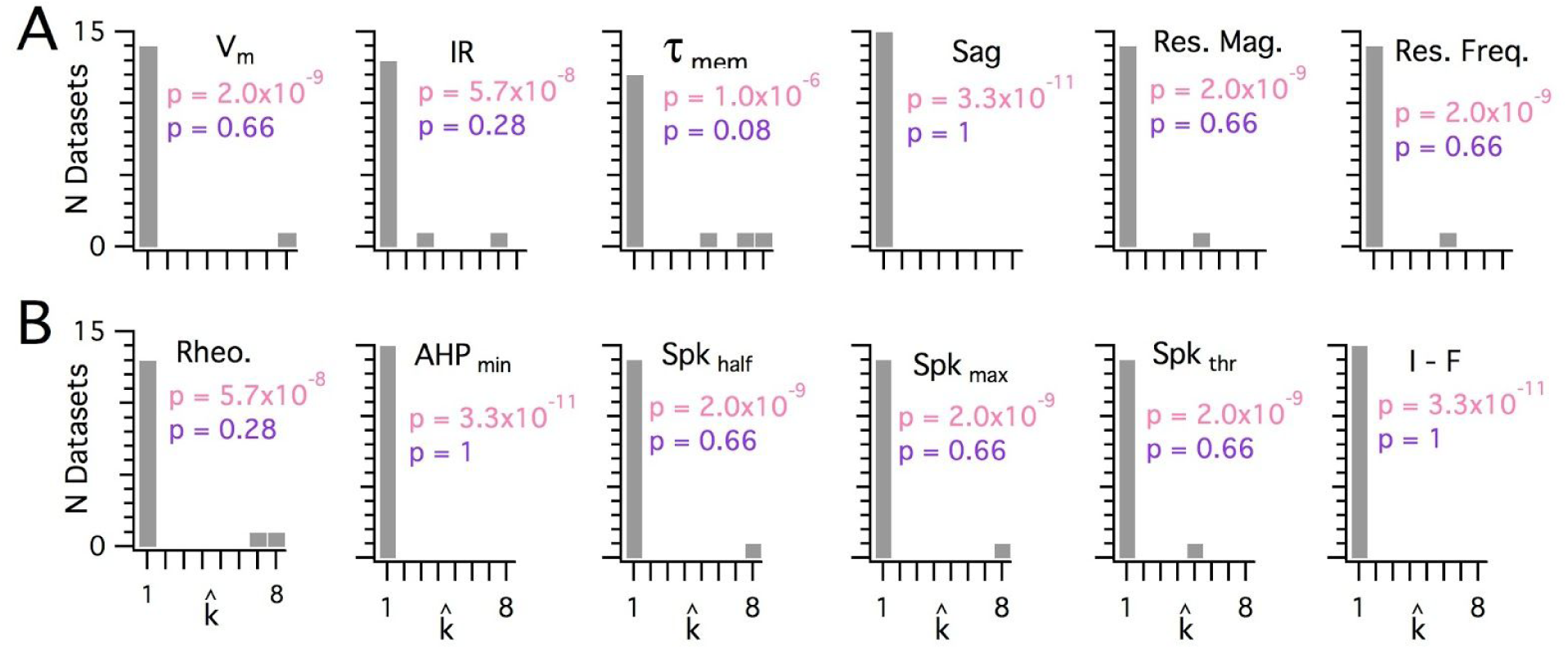
Significant modularity is not detectable for any measured property. (A-B) Histograms showing the k_est_ counts across all mice for each different measured sub-threshold (A) and supra-threshold (B) intrinsic properties. The maximum k evaluated was 8. The proportion of each measured property with k_est_ > 1 was compared using binomial tests (with N = 15) to the proportion expected if the underlying distribution of that property is always clustered with all separation between modes ≥ 5 std (pink text), or if the underlying distribution of the property is uniform (purple text). For all measured properties the values of k_est_ were indistinguishable (p > 0.05) from the data generated from a uniform distribution and differed from the data generated from a multi-modal distribution (p < 1 × 10^-6^).

When we investigated the other measured integrative properties we also failed to find evidence for modularity. Across all properties, for any given property at most 3 out of 15 mice were evaluated as having a clustered organisation using the modified gap statistic (Figure 6). This does not differ significantly from the proportion expected by chance when no modularity is present (p > 0.05, binomial test). Consistent with this, the average proportion of datasets evaluated as modular across all measured features was 0.072 ± 0.02 (± SEM), which is similar to the expected false positive rate. In contrast, for the 7 grid datasets with n ≥ 20 considered in Supplemental Figure 5, the mean for p_detect_ is 0.86, with 95 % confidence intervals of 0.42 to 0.996. We note here that previously published discontinuity algorithms did indicate significant modularity in the majority of the intrinsic properties measured in our dataset (N = 13/15 and N = 12/15 respectively), but this was attributable to false positives resulting from the relatively even sampling of recording locations (see Supplemental Figure 4A). Therefore, we conclude that it is unlikely that within individual animals any of the intrinsic properties we examined have organisation resembling the modular organisation of grid cells in behaving animals.

### Multiple sources of variance contribute to diversity in stellate cell intrinsic properties

Finally, because many of the measured electrophysiological features of SCs emerge from shared ionic mechanisms (Dodson et al., 2011; Garden et al., 2008; Pastoll et al., 2012a), we asked whether dorsoventral tuning reflects a single core mechanism and if inter-animal differences are specific to this mechanism or manifest more generally.

Estimation of conditional independence for measurements at the level of individual neurons (Figure 7A) or individual animals (Figure 7B) was consistent with the expectation that particular classes of membrane ion channels influence multiple electrophysiologically measured features. The first 5 dimensions of a principal components analysis (PCA) of all measured electrophysiological features accounted for almost 80% of the variance (Figure 7C). Examination of the rotations used to generate the principal components suggested relationships between individual features that are consistent with our evaluation of the conditional independence structure of the measured features (cf. Figure 7D and 7A). When we fit the principal components using mixed models with location as a fixed effect and animal identity as a random effect, we found that the first two components depended significantly on dorsoventral location (Figure 7E and Supplemental Table 14)(marginal R^2^ = 0.50 and 0.09 and adjusted p = 1.09 × 10^-15^ and 1.05 × 10^-4^ respectively). Thus, the dependence of multiple electrophysiological features on dorsoventral position may be reducible to two core mechanisms that together account for much of the variability between SCs in their intrinsic electrophysiology.

**Figure 7.**
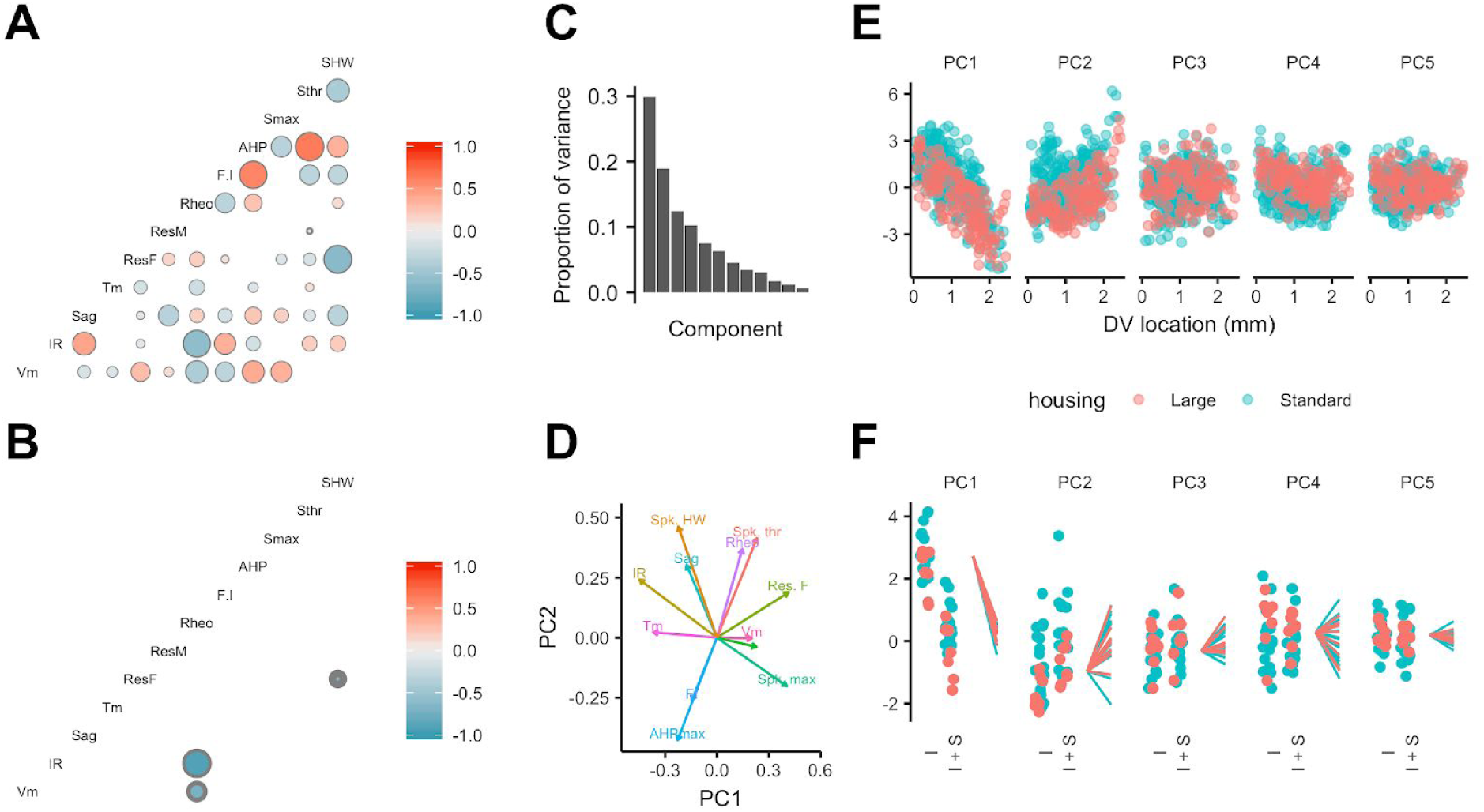
Feature relationships and inter-animal variability after reducing dimensionality of the data. (A-B) Partial correlations between the electrophysiological features investigated at the level of individual neurons (A) and at the level of animals (B). Partial correlations outside the 95% basic bootstrap confidence intervals are colour coded. Non-significant correlations are coloured white. (C) Proportion of variance explained by each principal component. To remove variance caused by animal age and the experimenter identity, the principal component analysis used a reduced dataset obtained by one experimenter and restricted to animals between 32 and 45 days old (N = 25, n = 572). (D) Loading plot for the first two principal components. (E) The first five principal components plotted as a function of position. (F) Intercept (I), intercept plus the slope (I + S) and slopes aligned to the same intercept, for fits for each animal of the first five principal components to a mixed model with location as a fixed effect and animal as a random effect.

Is inter-animal variation present in PCA dimensions that account for dorsoventral variation? The intercept, but not the slope of the dependence of the first two principal components on dorsoventral position depended on housing (adjusted p = 0.039 and 0.027)(Figure 7E, F and Supplemental Table 15). After accounting for housing, the first two principal components were still better fit by models that include animal identity as a random effect (adjusted p = 3.3 × 10^-9^ and 4.1 × 10^-86^), indicating remaining inter-animal differences in these components (Supplemental Table 16). A further 9 of the next 10 higher order principal components did not depend on housing (adjusted p > 0.1)(Supplemental Table 15), while 8 differed significantly between animals (adjusted p < 0.05)(Supplemental Table 16).

Together, these analyses indicate that dorsoventral organisation of multiple electrophysiological features of SCs reflect two main sources of variance, both of which are dependent on experience. Significant inter-animal variation in the major sources of variance remains after accounting for experience and experimental parameters.

## Discussion

Phenotypic variation is found across many areas of biology (Geiler-Samerotte et al., 2013), but has received little attention in investigations of mammalian nervous systems. We find unexpected inter-animal variability in SC properties suggesting that integrative properties of neurons are determined by set points that differ between animals and are controlled at a circuit level (Figure 8). Continuous, location-dependent organisation of set points for SC integrative properties provides new constraints on models for grid cell firing. More generally, the existence of inter-animal differences in set points has implications for experimental design and raises new questions about how integrative properties of neurons are specified.

**Figure 8.**
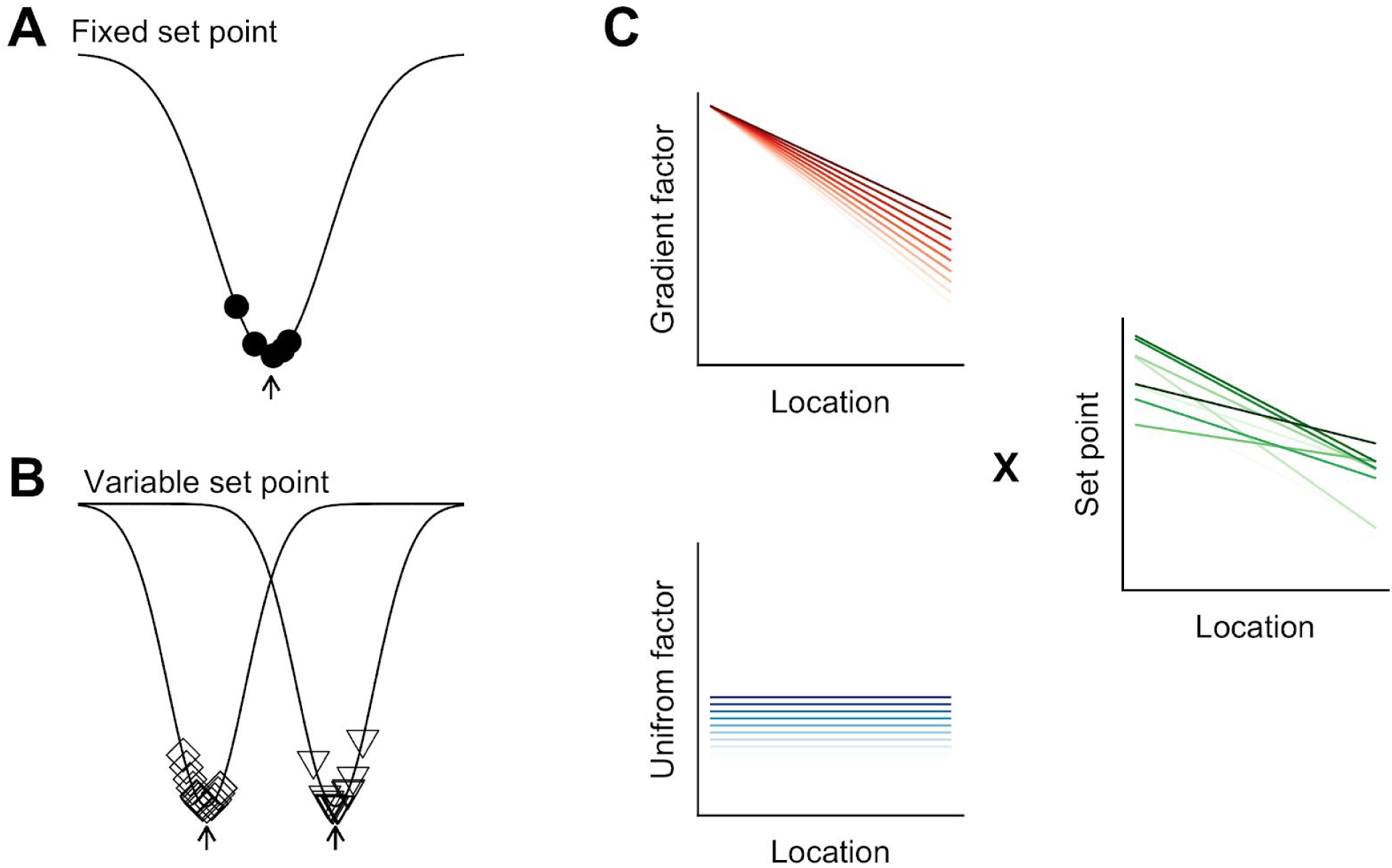
Models for intra- and inter-animal variation. (A) The configuration of a cell type can be conceived of as a trough (arrow head) in a developmental landscape (solid line). In this scheme, the trough can be considered as a set point. Differences between cells (filled circles) reflect variation away from the set point. (B) Neurons from different animals or located at different dorsoventral positions can be conceptualised as arising from different troughs in the developmental landscape. (C) Variation may reflect inter-animal differences in factors that establish gradients (upper left) and in factors that are uniformly distributed combing to generate set points that depend on animal identity and location. Each line corresponds to schematised properties of a single animal.

### A conceptual framework for within cell type variability

Theoretical models suggest how different cell types can be generated by varying target concentrations of intracellular Ca^2+^ or rates of ion channel expression (O’Leary et al., 2014). Within cell type variability predicted by these models arises from different initial conditions and may explain variability in our data between neurons from the same animal at the same dorsoventral location (Figure 8A). In contrast, the dependence of integrative properties on position and their variation between animals implies additional mechanisms that operate at circuit level (Figure 8B). In principle, this variation could be accounted for by inter-animal differences in dorsoventrally tuned or spatially uniform factors that influence ion channel expression or target points for intracellular Ca^2+^ (Figure 8C).

The mechanisms for within cell type variability suggested by our results may differ from inter-animal variation described in invertebrate nervous systems. Whereas in invertebrates inter-animal variability is between properties of individual identified neurons (Goaillard et al., 2009), in mammalian nervous systems neurons are not individually identifiable and the variation we describe here is at the level of populations. From a developmental perspective in which cell identity is considered as a trough in a state-landscape through which each cell moves (Wang et al., 2011b), variation in the population of neurons of the same type could be conceived as cell autonomous deviations from set points corresponding to the trough (Figure 8A). Our finding that variability among neurons of the same type manifests between as well as within animals, could be explained by differences between animals in the position of the trough or set point in the developmental landscape (Figure 8B).

### Implications of continuous dorsoventral organisation of stellate cell integrative properties for grid cell firing

Dorsoventral gradients in electrophysiological features of SCs have stimulated cellular models for the organisation of grid firing (Burgess, 2008; Giocomo and Hasselmo, 2008b; Grossberg and Pilly, 2012; O’Donnell and Nolan, 2011; Widloski and Fiete, 2014). Increases in spatial scale following deletion of HCN1 channels (Giocomo et al., 2011), which in part determine the dorsoventral organisation of SC integrative properties (Garden et al., 2008; Giocomo and Hasselmo, 2009), support a relationship between electrophysiological properties of SCs and grid cell spatial scales. Our data argue against models that explain this relationship through single cell computations (Burgess, 2008; Giocomo et al., 2007), as in this case modularity of integrative properties of SCs is required to generate modularity of grid firing. A continuous dorsoventral organisation of electrophysiological properties of SCs could support modular grid firing generated by self-organising maps (Grossberg and Pilly, 2012), or by synaptic learning mechanisms (Kropff and Treves, 2008; Urdapilleta et al., 2017). It is less clear how a continuous gradient would affect predictions of grid firing from continuous attractor network models (Shipston-Sharman et al., 2016; Widloski and Fiete, 2014), which can account for modularity by limiting synaptic interactions between modules.

The continuous variation of electrophysiological features of SCs suggested by our analysis is consistent with continuous gradients in gene expression along layer 2 of the MEC (Ramsden et al., 2015). It is also consistent with single cell RNA sequencing analysis of other brain areas, which indicates that while expression profiles for some cell types cluster around a point in feature space, others lie along a continuum (Cembrowski and Menon, 2018). It will be interesting in future to determine whether gene expression continua establish corresponding continua of electrophysiological features (cf. (Liss et al., 2001)).

### Functional consequences of within cell type inter-animal variability

What are the functional roles of inter-animal variability? In the crab stomatogastric ganglion, inter-animal variation correlates with circuit performance (Goaillard et al., 2009). Accordingly, variation in intrinsic properties of SCs might correlate with differences in grid firing (Domnisoru et al., 2013; Gu et al., 2018; Rowland et al., 2018; Schmidt-Hieber and Häusser, 2013) or behaviours that rely on SCs (Kitamura et al., 2014; Qin et al., 2018; Tennant et al., 2018). It is interesting in this respect that there appear to be inter-animal differences in the spatial scale of grid modules (cf. Figure 5 of (Stensola et al., 2012)). Modification of grid field scaling following deletion of HCN1 channels is also consistent with this possibility (Giocomo et al., 2011; Mallory et al., 2018). Alternatively, inter-animal differences may reflect multiple ways to achieve a common higher order phenotype. According to this view, coding of spatial location by SCs would not differ between animals despite lower level variation in their intrinsic electrophysiological features. This is related to the idea of degeneracy at the level of single cell electrophysiological properties (Marder and Goaillard, 2006; Mittal and Narayanan, 2018; O’Leary et al., 2014; Swensen and Bean, 2005), except that here the electrophysiological features differ between animals but higher order circuit computations may nevertheless be similar.

In conclusion, our results identify substantial within cell type variation in neuronal integrative properties that manifests between as well as within animals. This has implications for experimental design and model building as the distribution of replicates from the same animal will differ from those obtained from different animals (Marder and Taylor, 2011). An important future goal will be to distinguish causes of inter-animal variation. It is possible that additional external factors such as social interactions may affect brain circuitry (Wang et al., 2011a, 2014), although these effects appear focussed on frontal cortical structures rather than circuits for spatial computations (Wang et al., 2014). Alternatively, stochastic mechanisms operating at the population level may drive emergence of inter-animal differences during development of SCs (Donato et al., 2017; Ray and Brecht, 2016). Addressing these questions may turn out to be critical to understanding the relationship between cellular biophysics and circuit level computations in cognitive circuits (Schmidt-Hieber and Nolan, 2017).

## Acknowledgements

We thank Vanessa Stempel for comments on the manuscript, Tor Stensola and Edvard Moser for sharing published data, and Lukas Solanka and Lukas Fischer for help with building the large cage. This work was supported by grants to MN from the Wellcome Trust (200855/Z/16/Z) and the BBSRC (BB/L010496/1, BB/1022147/1 and BB/H020284/1).

## Author contributions

HP and MFN conceived of the study. HP and DG performed experiments. HP and MFN analysed the data. IP contributed to statistical analyses. GS assisted with experiments and contributed reagents. MFN obtained funding, supervised the study and wrote the paper. All authors contributed to review and editing of the manuscript.

## Methods

### Mouse strains

All experimental procedures were performed under a United Kingdom Home Office license and with approval of the University of Edinburgh’s animal welfare committee. Recordings of many SCs per animal used C57/Bl6J mice (Charles River). Recordings targeting calbindin cells used a Wfs1^Cre^ line (Wfs1-Tg3-CreERT2) obtained from Jackson Labs (Strain name: B6;C3-Tg(Wfs1-cre/ERT2)3Aibs/J; stock number:009103). Mice were group housed on a 12 h light/dark cycle (light on 07.30–19.30 h). Mice were usually housed in standard breeding cages, with a subset of mice instead housed from pre-weaning ages in a larger 2.4 × 1.2 m cage that was enriched with bright plastic objects (Supplemental Figure 6). All experiments in the standard cage used male mice. For experiments in the large cage, 2 mice were female, 6 mice were male and one was not identified. Because we did not find significant effects of sex on individual electrophysiologically properties all mice were included in the analyses reported in the text. When including only male mice the effects of housing on the first principal component remained significant, while the effects of housing on individual electrophysiologically properties no longer reach statistical significance after correcting for multiple comparisons. Additional analyses that consider only male mice are provided in the code associated with the manuscript.

### Viral injections

To label Cre expressing neurons in Wfs1^Cre^ heterozygous mice we stereotaxically injected AAV-FLEX-GFP, generated with a chimeric 1/2 serotype, which expresses GFP from a CBA promoter (Murray et al., 2011)(titer: 1.5 × 1012 cp/ml) as described in Surmeli et al., 2015.

### Slice preparation

Methods for preparation of parasagittal brain slices containing medial entorhinal cortex were based on procedures described previously (Pastoll et al., 2012b). Briefly, mice were sacrificed by cervical dislocation and their brains carefully removed and placed in cold (2-4 °C) modified ACSF, with composition (in mM): NaCl 86, NaH PO 1.2, KCl 2.5, NaHCO_3_ 25, Glucose 25, Sucrose 75, CaCl_2_ 0.5, MgCl_2_ 7. Brains were then hemisected and sectioned, also in modified ACSF at 4-8 °C, using a vibratome (Leica VT1200S). After cutting brains were placed at 36 °C for 30 minutes in standard ACSF, with composition (in mM): NaCl 124, NaH_2_PO4 1.2, KCl 2.5, NaHCO_3_ 25, Glucose 20, CaCl_2_ 2, MgCl_2_ 1. They were then allowed to cool passively to room temperature. All slices were prepared by the same experimenter (HP) following the same procedure on each day.

### Recording methods

Whole-cell patch-clamp recordings were made between 1 to 10 hours after slice preparation using procedures described previously (Pastoll et al., 2012a, 2012b, 2013; Sürmeli et al., 2015). Patch electrodes were filled with the following intracellular solution (in mM): K Gluconate 130; KCl 10, HEPES 10, MgCl_2_ 2, EGTA 0.1, Na_2_ATP 2, Na_2_GTP 0.3 NaPhosphocreatine 10. The open tip resistance was 4-5 MΩ, all seal resistances were > 2 GΩ and series resistance were < 30 MΩ. Recordings were made in current clamp mode using Multiclamp 700B amplifiers (Molecular Devices, Sunnyvale, CA, USA) connected to PCs via Instrutech ITC-18 interfaces (HEKA Elektronik, Lambrecht, Germany) and using Axograph X acquisition software (http://axographx.com/). Pipette capacitance and series resistances were compensated using the capacitance neutralisation and bridge-balance amplifier controls. An experimentally measured liquid junction potential of 12.9 mV was not corrected for. Stellate cells were identified by their large sag response and the characteristic waveform of their action potential afterhyperpolarization (see (Alonso and Klink, 1993; Gonzalez-Sulser et al., 2014; Nolan et al., 2007; Pastoll et al., 2012a)).

To maximize the number of cells recorded per animal we adopted two strategies. First, to minimize the time required to obtain data from each recorded cell, we measured electrophysiological features using a series of three short protocols following initiation of stable whole-cell recordings. We used responses to sub-threshold current steps to estimate passive membrane properties (Figure 2A), a frequency modulated sinusoidal current waveform (ZAP waveform) to estimate impedance amplitude profiles (Figure 2B) and a linear current ramp to estimate the action potential threshold and firing properties (Figure 2C). From analysis of data obtained with these protocols we obtained 12 quantitative measures that describe sub- and supra-threshold integrative properties of each recorded cell (Figure 2A-C). Second, for the majority of mice two experimenters recorded in parallel from neurons in two sagittal brain sections from the same hemisphere. The median dorsal-ventral extent of the recorded SCs was 1644 µm (range 0 - 2464 µm). Each experimenter aimed to evenly sample recording locations across the dorsoventral extent of the MEC, and for most animals each experimenter recorded sequentially from opposite extremes of the dorsoventral axis. Each experimenter varied the starting location for recording between animals. For several features the direction of recording affected their measured dependence on dorsoventral location, but the overall dependence of these features on dorsoventral location was robust to this effect (Supplemental Table 9).

### Measurement of electrophysiological features

Electrophysiological recordings were analysed in Matlab (Mathworks) using a custom written semi-automated pipeline. We defined each feature as follows (see also (Nolan et al., 2007; Pastoll et al., 2012a)): resting membrane potential was the mean of the membrane potential during 1 s prior to injection of current steps used to assess subthreshold properties; input resistance was the steady-state voltage response to the negative current steps divided by their amplitude; membrane time constant was the time constant of an exponential function fit to the initial phase of membrane potential responses to the negative current steps; the sag coefficient was the steady-state divided by the peak membrane potential response to the negative current steps; resonance frequency was the frequency at which the peak membrane potential impedance was found to occur; resonance magnitude was the ratio between the peak impedance and the impedance at a frequency of 1 Hz; action potential threshold was calculated from responses to positive current ramps as the membrane potential at which the first derivative of the membrane potential crossed 20 mv ms^-1^ averaged across the first five spikes, or fewer if less spikes were triggered; rheobase was the ramp current at the point when the threshold was crossed on the first spike; spike maximum was the mean peak amplitude of the action potentials triggered by the positive current ramp; spike width was the duration of the action potentials measured at the voltage corresponding to the midpoint between the spike threshold and spike maximum; the AHP minimum was the negative peak membrane potential of the afterhyperpolarization following the first action potential when a second action potential also occurred; the F-I slope was the linear slope of the relationship between the spike rate and injected ramp current over a 500 ms window.

### Analysis of location-dependence, experience-dependence and inter-animal differences

Analyses of location-dependence and inter-animal differences used R 3.4.3 (R Core Team, Vienna, Austria) and R Studio 1.1.383 (R Studio Inc., Boston, MA).

To fit linear mixed effect models we used the lme4 package (Bates et al., 2014). Features of interest were included as fixed effects and animal identity was included as a random effect. All reported analyses are for models with the minimal a priori random effect structure, in other words the random effect was specified with uncorrelated slope and intercept. We also evaluated models in which only the intercept, or correlated intercept and slope were specified as the random effect. Model assessment was performed using Akaike Information Criterion (AIC) scores. In general, models with either random slope and intercept, or correlated random slope and intercept, had lower AIC scores than random intercept only models, indicating a better fit to the data. We used the former set of models for all analyses of all properties for consistency and because a maximal effect structure may be preferable on theoretical grounds (Barr et al., 2013). We evaluated assumptions including linearity, normality, homoscedasticity and influential data points. For some features we found modest deviations from these assumptions that could be remedied by handling non-linearity in the data using copula transformation. Model fits were similar following transformation of the data. However, we focus here on analyses of the untransformed data because physical interpretation of values for data points is clearer.

To evaluate location-dependence of a given feature, p-values were calculated using a *χ*^2^ test comparing models to null models with no location information but identical random effect specification. To calculate marginal and conditional R^2^ of mixed effect models we used the MuMin package (Bartoń, 2014). To evaluate additional fixed effects we used Type II Wald *χ*^2^ test tests provided by the car package (Fox and Weisberg, 2018). To compare mixed effect with equivalent linear models we used a *χ*^2^ test to compare the calculated deviance for each model. For clarity we have reported key statistics in the main text and provide full test statistics in the Supplemental Tables. In addition the code will be provided as an R markdown document in which the analyses can be fully reproduced.

To evaluate partial correlations between features we used the function cor2pcor from the R package corpcor (Schafer et al., 2017). Principal components analysis used core R functions.

### Implementation of tests for modularity

To establish statistical tests to distinguish ‘modular’ from ‘continuous’ distributions given relatively few observations, we classified datasets as continuous or modular by modifying the gap statistic algorithm (Tibshirani et al., 2001). The gap statistic estimates the number of clusters (k_est_) that best account for the data in any given dataset (Supplementary Figure 1A-C). However, this estimate may be prone to false positives, particularly where the numbers of observations are low. We therefore introduced a thresholding mechanism for tuning the sensitivity of the algorithm so that the false positive rate, which is the rate of misclassifying datasets drawn from continuous (uniform) distributions as ‘modular’, is low, constant across different numbers of cluster modes and insensitive to dataset size (Supplementary Figure 1D-G). With this approach we are able to estimate whether a dataset is best described as lacking modularity (k_est_ = 1), or having a given number of modes (k_est_ > 1). We describe below tests carried out to validate the approach.

To illustrate the sensitivity and specificity of the modified gap statistic algorithm we applied it to simulated datasets drawn either from a uniform distribution (k = 1, n = 40) or a bimodal distribution with separation between the modes of 5 standard deviations (k = 2, n = 40, sigma = 5)(Supplemental Figure 2A). We set the thresholding mechanism so that k_est_ for each distinct k (where k_est_ ≥ 2) has a false positive rate of 0.01. In line with this, testing for 2 ≤ k_est_ ≤ 8 (the maximum k expected to occur in grid spacing in the MEC), across multiple (N = 1000) simulated datasets drawn from the uniform distribution, produced a low false positive rate (P(k_est_) ≥ 2 = ∼0.07), whereas when the data were drawn from the bimodal distribution the ability to detect modular organisation (p_detect_) was good (P(k_est_) ≥ 2 = ∼ 0.8)(Supplemental Figure 2B). The performance of the statistic improved with larger separation between clusters and with greater numbers of data points per dataset (Supplemental Figure 2C) and is relatively insensitive to the numbers of clusters (Supplemental Figure 2D). The algorithm maintains high rates of p_detect_ when modes have varying densities and when sigma between modes varies in a manner similar to grid spacing data (Supplemental Figure 3).

Recently described algorithms (Giocomo et al., 2014; Stensola et al., 2012) address the problem of identifying modularity when data are sampled from multiple locations and data values vary as a function of location, as is the case for the mean spacing of grid fields for cells at different dorsoventral locations (Hafting et al., 2005). They generate log normalised discontinuity (which we refer to here as lnDS) or discreteness scores, which are the log of the ratio of discontinuity or discreteness scores for the data points of interest and for the sampling locations, with positive values interpreted as evidence for clustering (Giocomo et al., 2014; Stensola et al., 2012). However, in simulations of datasets generated from a uniform distribution with evenly spaced recording locations, we find that the lnDS is always greater than zero (Supplemental Figure 4A). This is because evenly spaced locations result in a discontinuity score that approaches zero and therefore the log ratio of the discontinuity of the data to this score will be positive. Thus, for evenly spaced data the lnDS is guaranteed to produce false positive results. When locations are instead sampled from a uniform distribution, approximately a half of simulated datasets have a log discontinuity ratio greater than 0 (Supplemental Figure 4A), which in previous studies would be interpreted as evidence for modularity (Giocomo et al., 2014). Similar discrepancies arise for the discreteness measure (Stensola et al., 2012). To address these issues we introduced a log discontinuity ratio threshold, so that the discontinuity method could be matched to produce a similar false positive rate to the adapted gap statistic algorithm used in the example above. After including this modification, we found that for a given false positive rate the adapted gap statistic is more sensitive at detecting modularity in the simulated datasets (Supplementary Figure 4B).

To establish whether the modified gap statistic detects clustering in experimental data we applied it to previously published grid cell data (Stensola et al., 2012). We find that the modified gap statistic identified clustered grid spacing for 6 of 7 animals previously identified as having grid modules and with n ≥ 20. For these animals the number of modules was similar (but not always identical) to the number of previously identified modules (Supplementary Figure 5). In contrast, the modified gap statistic does not identify clustering in 5 of 6 sets of recording locations, confirming that the grid clustering is likely not a result of uneven sampling of locations (we could not test the 7^th^ as location data were not available). The thresholded discontinuity score also detects clustering in the same 5 of the 6 tested sets of grid data. From the 6 grid datasets detected as clustered with the modified gap statistic we estimated the separation between clusters by fitting the data with a mixture of Gaussians, with the number of modes set by the value k obtained with the modified gap statistic. This analysis suggested that the largest spacing between contiguous modules in each mouse is always > 5.6 standard deviations (mean = 20.5 ± 5.0 standard deviations). Thus, the modified gap statistic detects modularity found in the grid system and indicates that previous descriptions of grid modularity are in general robust to the possibility of false positives associated with the discreteness and discontinuity methods.

## Data and code availability

Data will be made available through the University of Edinburgh Datashare resource (https://datashare.is.ed.ac.uk/). Code for analyses is available at https://github.com/MattNolanLab/Inter_Intra_Variation.

---

**Supplemental Figure 1.**
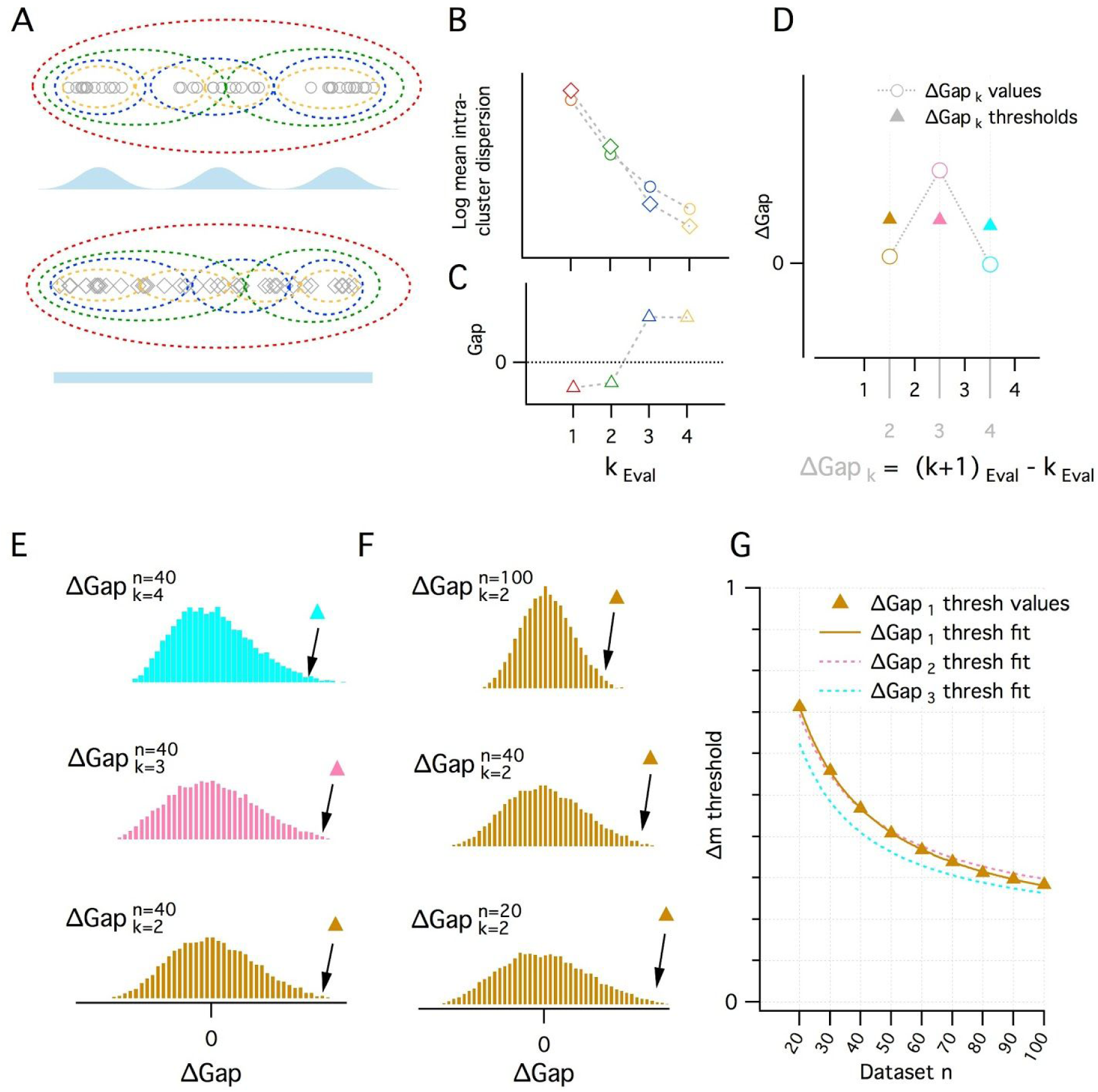
A quantitative adaptation of the Gap Statistic clustering algorithm. (A-C) Logic of the Gap Statistic. (A) Simulated clustered dataset with three modes (k = 3)(open grey circles) (upper) and corresponding simulated reference dataset drawn from a uniform distribution with lower and upper limits set by the minimum and maximum values from the corresponding clustered dataset (open grey diamonds). Data were allocated to clusters by running a K-means algorithm 20 times on each set of data and selecting the partition with the lowest average intracluster dispersion. Red, green, blue and yellow dashed ovals show a realisation of data subsets allocated to each cluster when k_Eval_ = 1, 2, 3 and 4 modes. (B) The average value of the log intracluster dispersion for the clustered (open circles) and reference (open diamonds) datasets for each value of k tested in (A). (C) Gap values resulting from the difference between the clustered and reference values for each k in (B). Many (≥ 500 in this paper) reference distributions are generated and their mean intracluster dispersion values are subtracted from those arising from the clustered distribution to maximise the reliability of Gap values. (D) A procedure for selecting the optimal k depending on the associated gap values. Quantitative procedure for selecting optimal k (k_est_). ΔGap values (open circles) are obtained by subtracting from the Gap value of a given k the Gap value for the previous k (ΔGap_k_ = Gap_k_ - Gap_k-1_). For each ΔGap_k_, if its ΔGap value greater than a threshold (filled triangles) associated with that ΔGap_k_, that ΔGap_k_ will be k_est_, if no ΔGap exceeds its threshold, k_est_ = 1. In the figure, for ΔGap_k_ = 2, 3, 4 (brown, pink and cyan), the ΔGap value exceeds its threshold only when ΔGap_k_ = 3. Therefore k_est_ = 3. (E-G) Determination of ΔGap_k_ thresholds. (E) Histograms of ΔGap values calculated from 20000 simulated datasets drawn from uniform distributions for each ΔGap_k_ (brown, pink and cyan respectively for ΔGap_k_ = 2, 3, 4) for a single dataset size (n = 40). ΔGap thresholds (filled triangles) are the values beyond which 1% of the ΔGap values fall and vary with ΔGap_k_. (F) Histograms of ΔGap values for a range of dataset sizes (n = 20, 40, 100) and their associated thresholds. (G) Plot of the ΔGap thresholds as a function of dataset size and ΔGap_k_. For separate ΔGap_k_, ΔGap threshold values are well fit by a hyperbolic function of dataset size. These fits provide a practical method of finding the appropriate ΔGap threshold for an arbitrary dataset size.

**Supplemental Figure 2.**
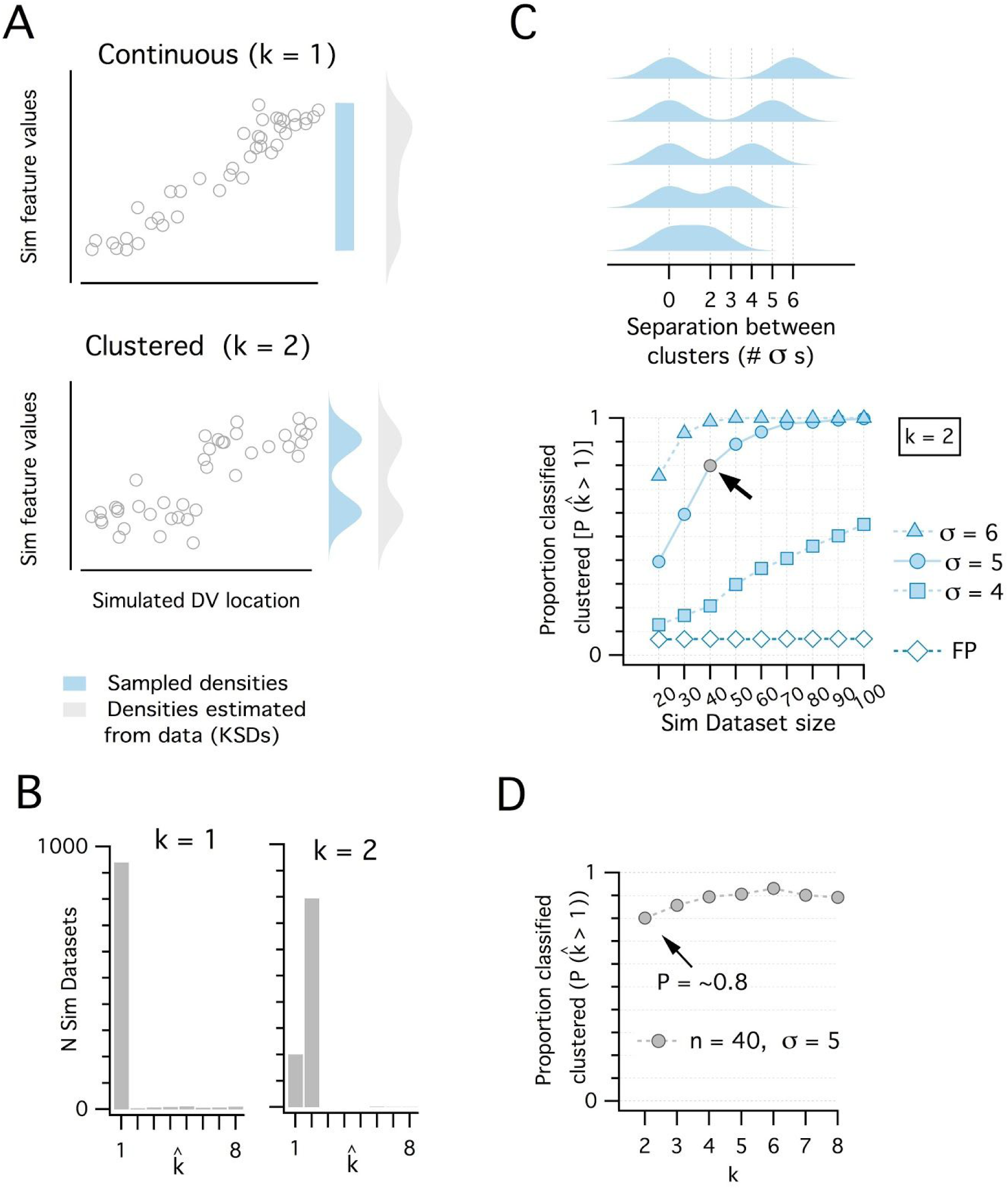
Discrimination of continuous from modular organisations using the adapted Gap Statistic algorithm. (A) Simulated datasets (each n = 40) drawn from continuous (uniform, k = 1 mode) (upper) and modular (multimodal mixture of Gaussians with k = 2 modes)(lower) distributions, and plotted against simulated dorsoventral locations. Also shown are probability density functions (pdf) used to generate each dataset (light blue) and the densities estimated post-hoc from the generated data as kernel smoothed densities (light grey pdfs). (B) Histograms showing the distribution of k_est_ from 1000 simulated datasets drawn from each pdf in (A). k_est_ is determined for each dataset by a modified gap statistic algorithm (see Supplemental Figure 1 above). When k_est_ = 1, the dataset is considered continuous (unclustered), when k_est_ ≥ 2 the dataset is considered modular (clustered). The algorithm operates only on the feature values and does not use location information. (C) Illustration of a set of clustered (k = 2) pdfs with the distance (in standard deviations) between clusters ranging from 2 to 6 (upper). Systematic evaluation of the ability of the modified gap statistic algorithm to detect clustered organisation (k_est_ ≥ 2) in simulated datasets of different size (n = 20 to 100) drawn from the clustered (filled blue) and continuous (open blue) pdfs (lower). The proportion of datasets drawn from the continuous distribution that have k_est_ ≥ 2 is the false positive (FP) rate (pFP = ∼0.07). The light grey filled circle shows the smallest dataset size (n = 40) with SD = 5 where the proportion of datasets detected as clustered (p_detect_) is ∼ 0.8. (D). Plot showing how p_detect_ at n = 40, SD = 5 changes when datasets are drawn from pdfs with different numbers of clusters (n modes from 2 to 8). Further evaluation of analysis of additional clusters is in the following figure.

**Supplemental Figure 3.**
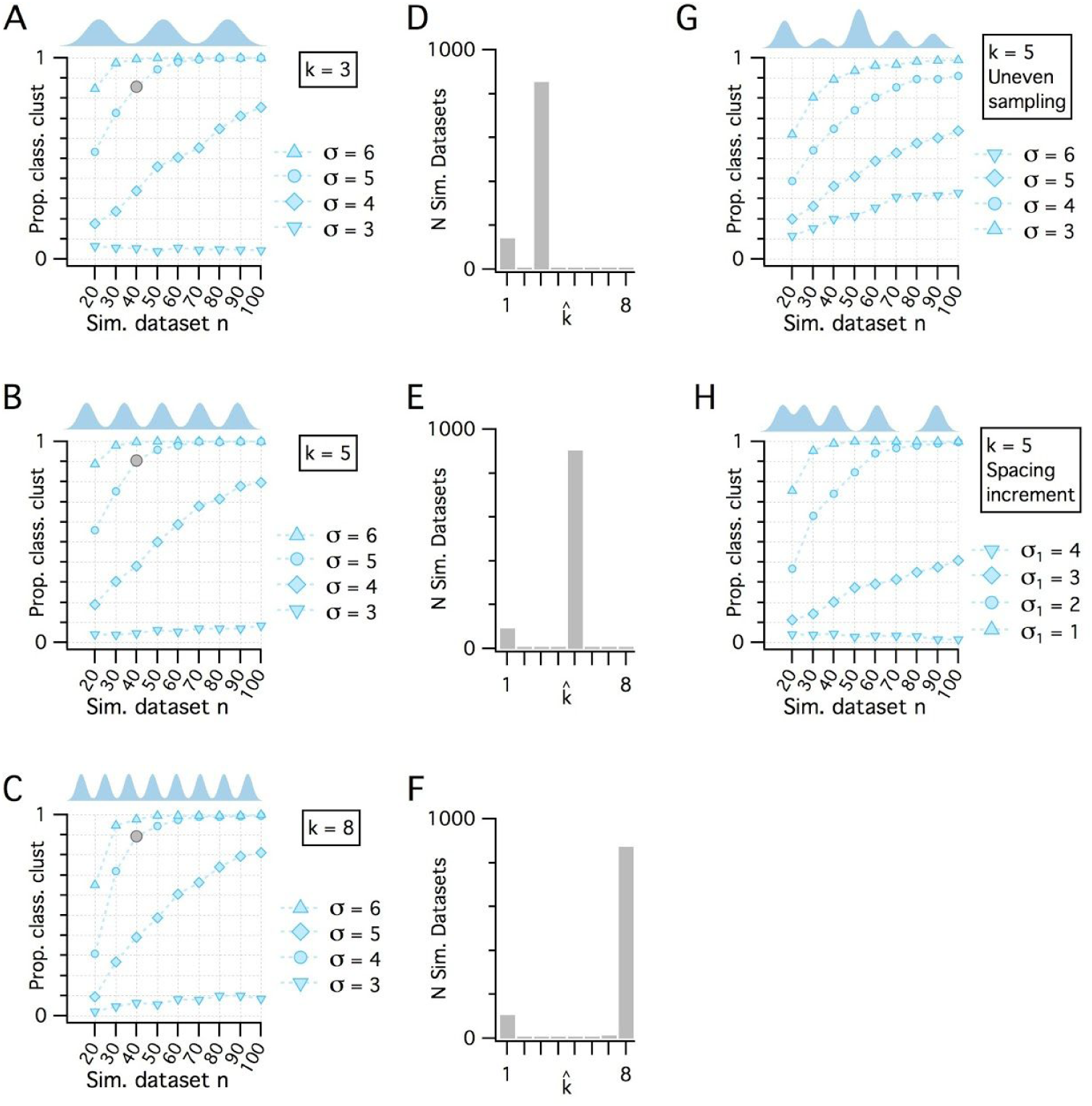
Additional evaluation of the adapted Gap Statistic algorithm. (A – C) Plots showing the results showing how p_detect_ (the ability of the modified gap statistic algorithm to detect clustered organisation) depends on dataset size and separation between cluster modes in simulated datasets drawn from clustered pdfs with different numbers of modes. The grey markers indicate n = 40, SD = 5 (as shown in Figure 1E). In each plot, p_detect_ is shown as a function of simulated dataset size and separation between modes when k = 3 (A), k = 5 (B) and k = 8 (C), which was the maximum number of clusters evaluated. (D – F) Histograms showing the counts of k_est_ from the 1000 simulated n = 40, SD = 5 datasets (grey filled circles) illustrated in (A) – (C) respectively. (G) p_detect_ as a function of dataset size and mode separation with k = 5 but when cluster modes are unevenly sampled. Sample sizes from clusters vary randomly with each dataset. A single mode can contribute from all to none of the points in any simulated dataset. (H) p_detect_ as a function of dataset size and mode separation with k = 5 but when the distance between mode centres increases by a factor of sqrt(2) between sequential cluster pairs. Data is shown for different initial separations (the distance between the first two cluster centres) ranging from 1 to 4 (with corresponding separations between the final cluster pair ranging from 4 to 16).

**Supplemental Figure 4.**
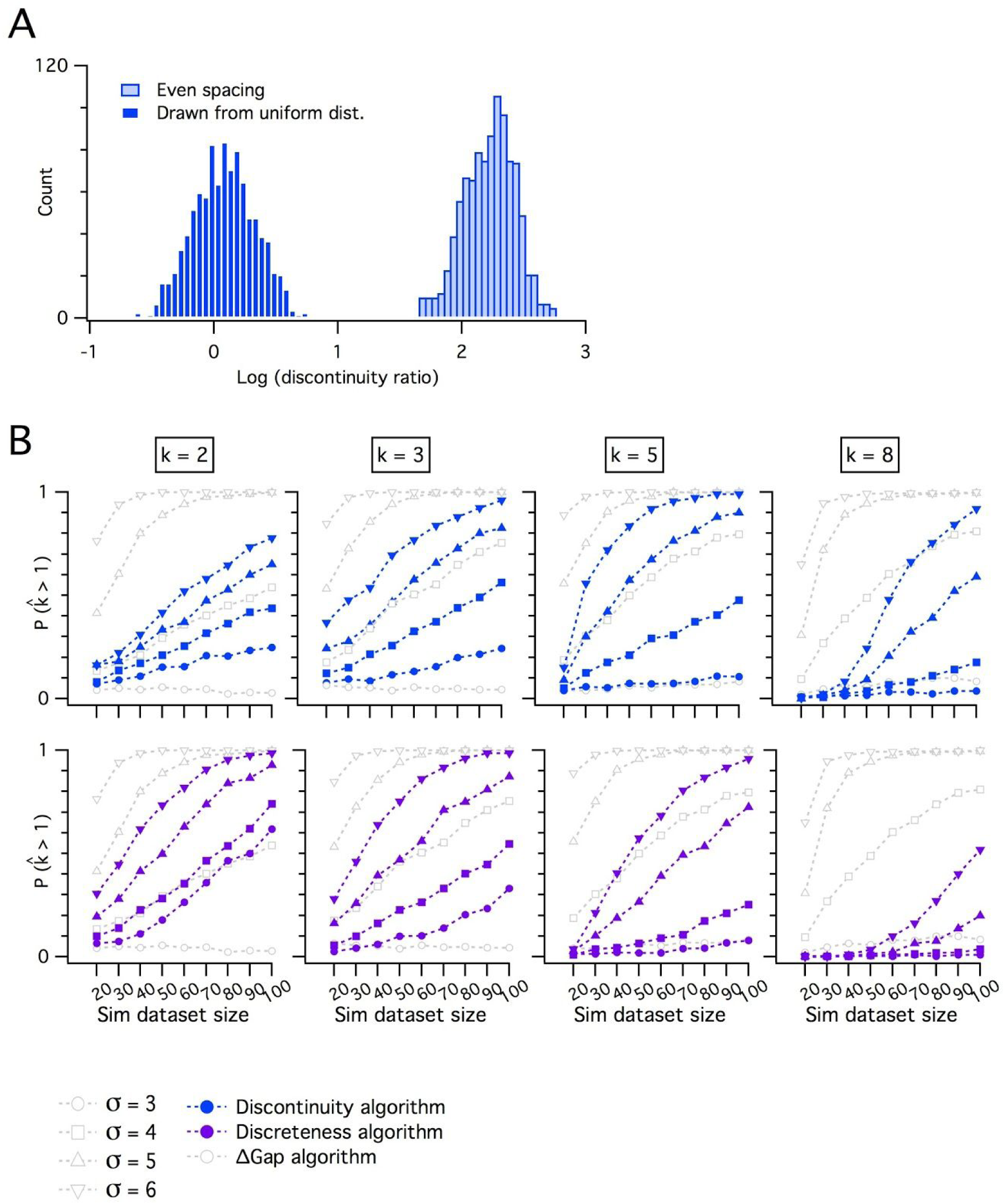
Comparing delta gap with discontinuity measures for discreteness. (A) Counts of log discontinuity ratio scores generated from a simulated uniform data distribution. The data distribution was ordered and then sampled either at positions drawn at random from a uniform distribution (dark blue), or at positions with a fixed increment (light blue). For the data sampled at random positions approximately half of the scores are > 0 and for even sampling all scores are > 0. Therefore, a threshold score > 0 does not distinguish discrete from continuous distributions. (B) Comparison of p_detect_ as a function of dataset size for the adapted Gap Statistic Algorithm, the discontinuity (upper) and the discreteness algorithm (lower). Each algorithm is adjusted to yield a 7% false positive rate. Each column shows simulations of data with different numbers of modes (k).

**Supplemental Figure 5.**
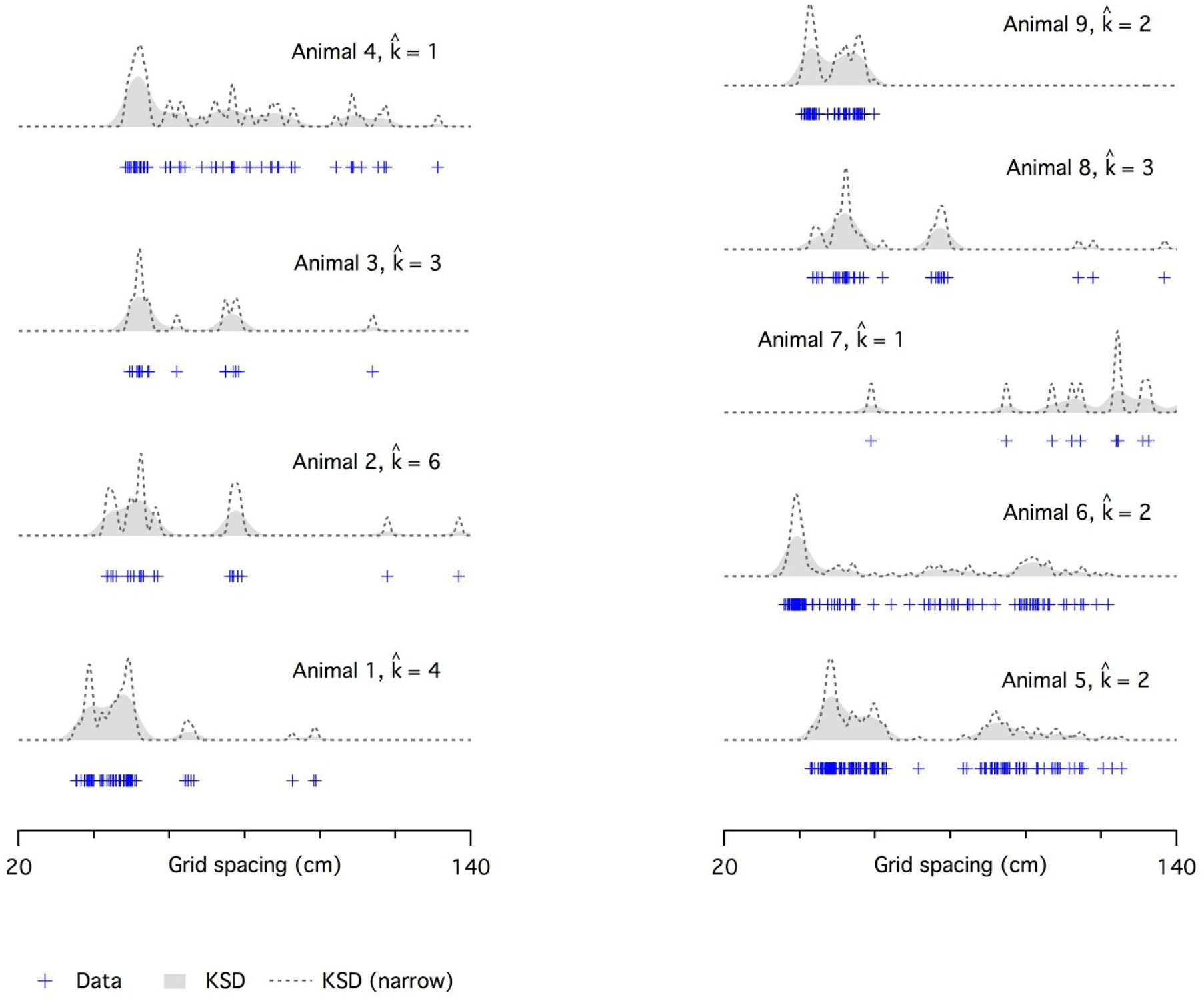
Evaluation of modularity of grid firing using a modified gap statistic algorithm. Examples of grid spacing for individual neurons (crosses) from different mice. Kernel smoothed densities (KSDs) were generated with either a wide (solid grey) or narrow (dashed lines) kernel. The number of modes estimated using the modified gap statistic algorithm ranges is ≥ 2 for all but one animal (animal 4) with n ≥ 20 (animals 3 and 7 have < 20 recorded cells). We did not have location information for animal 2.

**Supplemental Figure 6.**
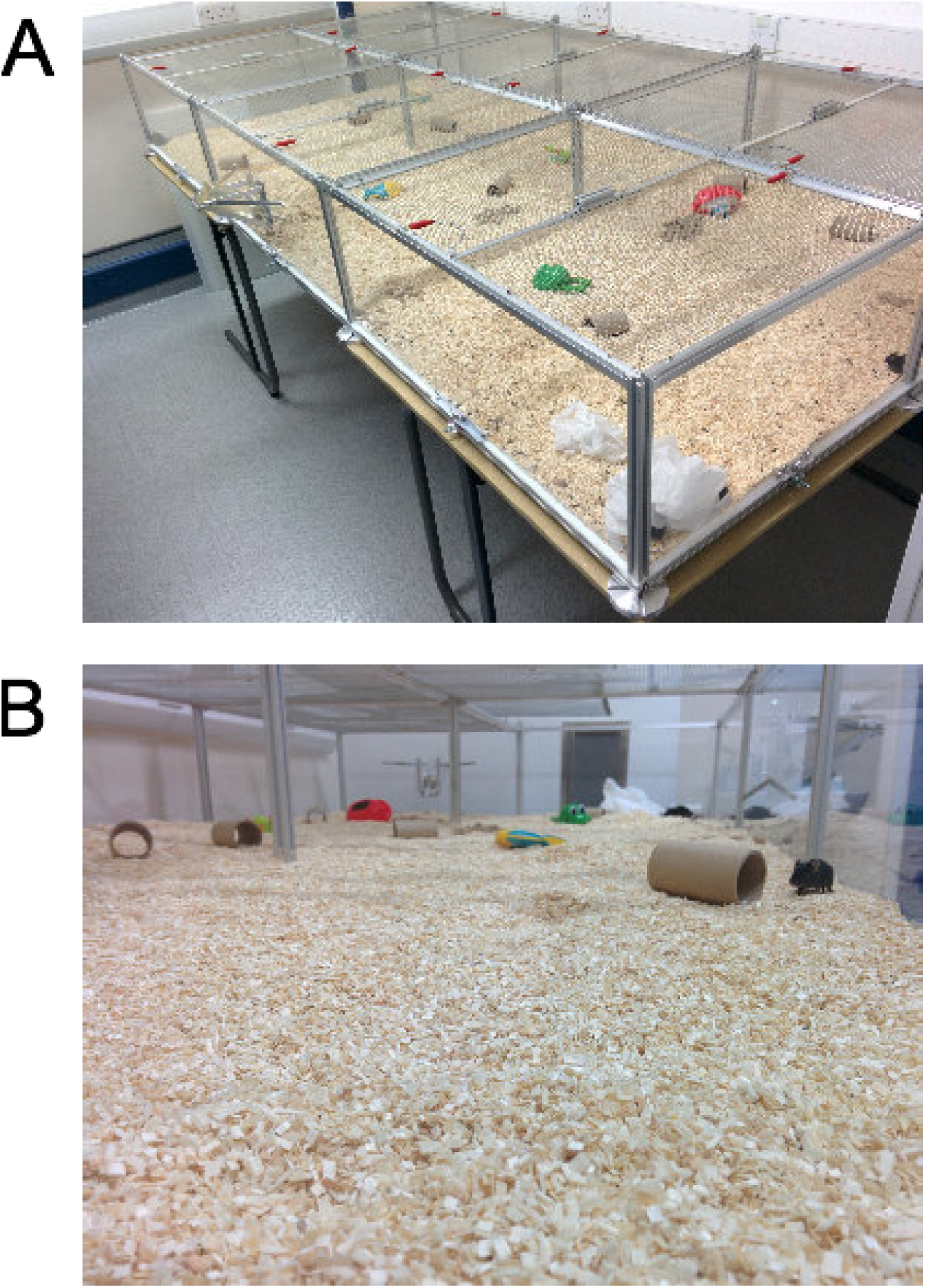
Large environment for housing. (A-B) The large cage environment viewed from above (A) and from inside (B).

**Supplemental Table 1.** Dependence of calbindin cell properties on dorsoventral position. Analyses are as described for Table 1. Data are from GFP positive putative pyramidal neurons (n = 42, N = 3).

| Property | Slope | p (slope) | Marginal R2 | Conditional R2 | Slope (min) | Slope (max) | p (vs linear) |
| --- | --- | --- | --- | --- | --- | --- | --- |
| Vm | 2.0086 | 0.1558 | 0.1104 | 0.1104 | 2.0086 | 2.0086 | 1 |
| IR | 0.1655 | 0.9652 | 0.0001 | 0.0181 | 0.1655 | 0.1655 | 1 |
| Sag | -0.0137 | 0.5928 | 0.0142 | 0.0142 | -0.0137 | -0.0137 | 1 |
| Tm | 3.0716 | 0.2531 | 0.0861 | 0.1681 | 1.7960 | 4.2054 | 1 |
| Res. frequency | 0.2200 | 0.8135 | 0.0042 | 0.0042 | 0.2200 | 0.2200 | 1 |
| Res. magnitude | 0.0940 | 0.3552 | 0.0673 | 0.1764 | 0.0623 | 0.1553 | 1 |
| Spike threshold | 2.5595 | 0.1558 | 0.0986 | 0.0986 | 2.5595 | 2.5595 | 1 |
| Spike maximum | 1.1065 | 0.4628 | 0.0280 | 0.1533 | 1.1065 | 1.1065 | 1 |
| Spike width | 0.0495 | 0.0298 | 0.1922 | 0.1922 | 0.0495 | 0.0495 | 1 |
| Rheobase | -15.4864 | 0.5928 | 0.0165 | 0.0165 | -15.4864 | -15.4864 | 1 |
| Spike AHP | 1.6079 | 0.3742 | 0.0470 | 0.0982 | 1.0252 | 2.2523 | 1 |
| I-F slope | -0.0037 | 0.8625 | 0.0019 | 0.0019 | -0.0037 | -0.0037 | 1 |

**Supplemental Table 2.**
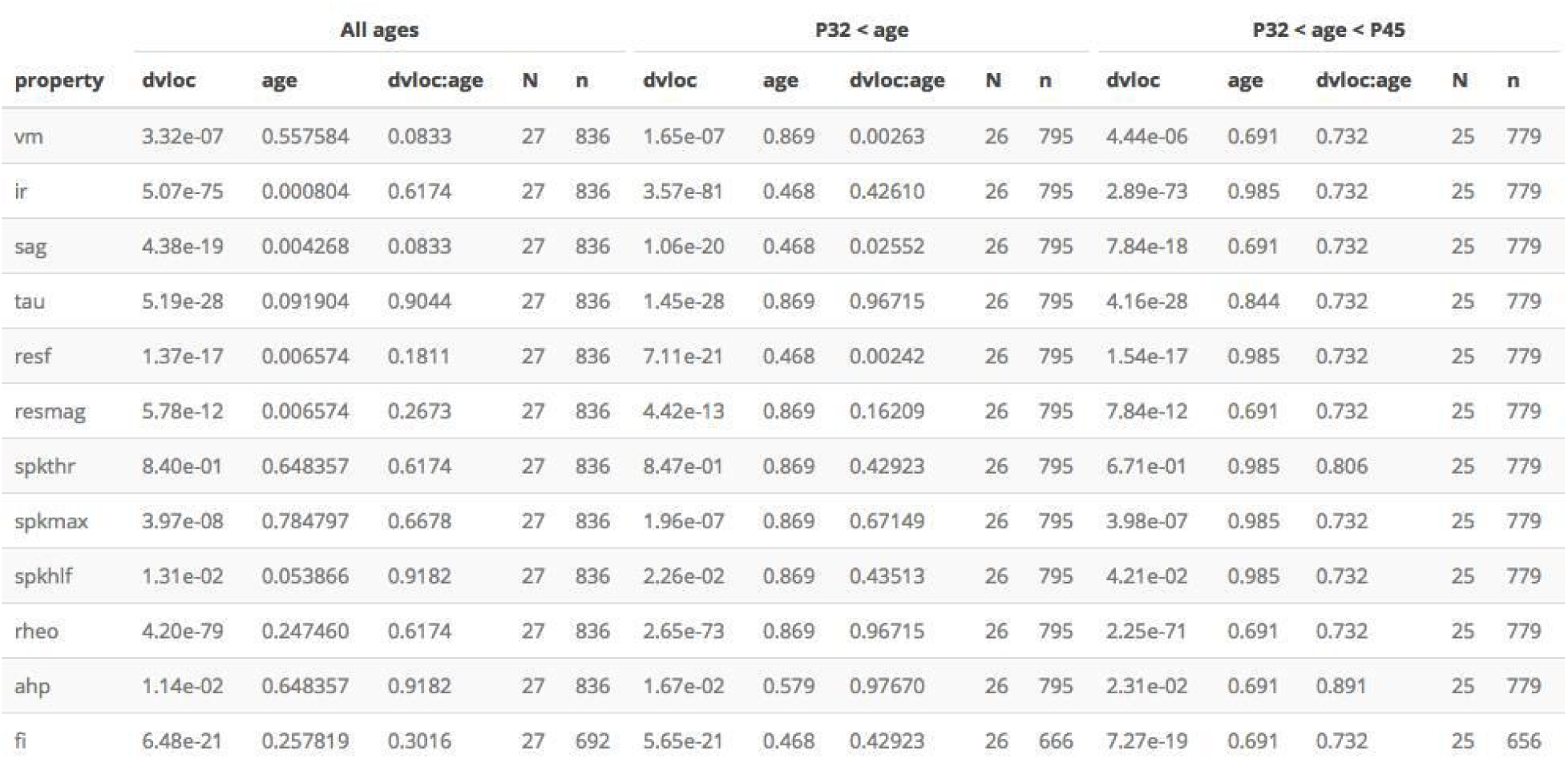
Dependence of SC properties on age. The distinguishing electrophysiological features of SCs and their dorsoventral organisation were apparent at all ages, with some features depending significantly on age (left columns), consistent with the idea that SCs continue to mature beyond P18 (Boehlen et al., 2010; Burton et al., 2008). When we considered only animals between P33 and P44 we did not find any significant effect of age (right columns). Significance estimates for the effects of dorsoventral position (dvloc), age (age) and interactions between dorsoventral position and age (dvloc:age) were estimated using type II ANOVA and Wald *χ*^2^ test from fits to mixed models containing age and location as fixed effects and animal identity as random effects. Significance estimates were adjusted for multiple comparisons using the Benjamini and Hochberg method.

**Supplemental Table 3.** Dependence of SC properties on housing. Significance estimates for the effects of dorsoventral position (dvloc), housing (housing) and interactions between dorsoventral position and housing (dvloc:housing) estimated using type II ANOVA and Wald *χ*^2^ test from fits to mixed models containing age and location as fixed effects and animal identity as random effects. Initial significance estimates (raw p) were adjusted for multiple comparisons (adjusted p) using the Benjamini and Hochberg method.

| property | N | n | Int | Fixed effects |  |  | raw p |  |  | adjusted p |  |  |
| --- | --- | --- | --- | --- | --- | --- | --- | --- | --- | --- | --- | --- |
|  |  |  |  | dvloc | housing | dv:housing | dvloc | housing | dv:housing | dvloc_adj | housing_adj | dv:housing_adj |
| vm | 25 | 779 | -63.657 | -0.836 | 0.276 | 0.0524 | 6.1e-06 | 0.5330 | 0.8873 | 8.2e-06 | 0.581 | 0.972 |
| ir | 25 | 779 | 20.015 | 11.288 | -2.306 | 0.3679 | 7.2e-74 | 0.0942 | 0.7807 | 4.3e-73 | 0.188 | 0.972 |
| sag | 25 | 779 | 0.531 | 0.032 | 0.023 | 0.0009 | 3.0e-19 | 0.0031 | 0.9042 | 8.9e-19 | 0.031 | 0.972 |
| tau | 25 | 779 | 8.408 | 2.301 | -1.110 | 0.4297 | 9.1e-28 | 0.0606 | 0.3824 | 3.7e-27 | 0.145 | 0.765 |
| resf | 25 | 779 | 9.187 | -1.024 | 1.189 | -0.2756 | 3.9e-17 | 0.0113 | 0.3543 | 7.8e-17 | 0.045 | 0.765 |
| resmag | 25 | 779 | 1.804 | -0.091 | 0.048 | -0.0339 | 8.7e-12 | 0.7457 | 0.3255 | 1.5e-11 | 0.746 | 0.765 |
| spkthr | 25 | 779 | -39.222 | 0.140 | 0.715 | 0.0275 | 6.8e-01 | 0.4332 | 0.9725 | 6.8e-01 | 0.537 | 0.972 |
| spkmax | 25 | 779 | 43.696 | 2.343 | 1.073 | -0.6996 | 3.3e-07 | 0.4478 | 0.3664 | 5.0e-07 | 0.537 | 0.765 |
| spkhlf | 25 | 779 | 0.501 | 0.022 | -0.055 | -0.0136 | 4.0e-02 | 0.0052 | 0.3060 | 4.4e-02 | 0.031 | 0.765 |
| rheo | 25 | 779 | 398.932 | -97.192 | 65.598 | -35.5348 | 7.3e-96 | 0.0275 | 0.0031 | 8.8e-95 | 0.083 | 0.038 |
| ahp | 25 | 779 | -56.474 | -0.470 | 0.980 | -0.2299 | 2.1e-02 | 0.2537 | 0.6794 | 2.5e-02 | 0.380 | 0.972 |
| fi | 25 | 656 | 0.034 | 0.039 | 0.011 | -0.0032 | 5.0e-19 | 0.1874 | 0.7107 | 1.2e-18 | 0.321 | 0.972 |

**Supplemental Table 4.**
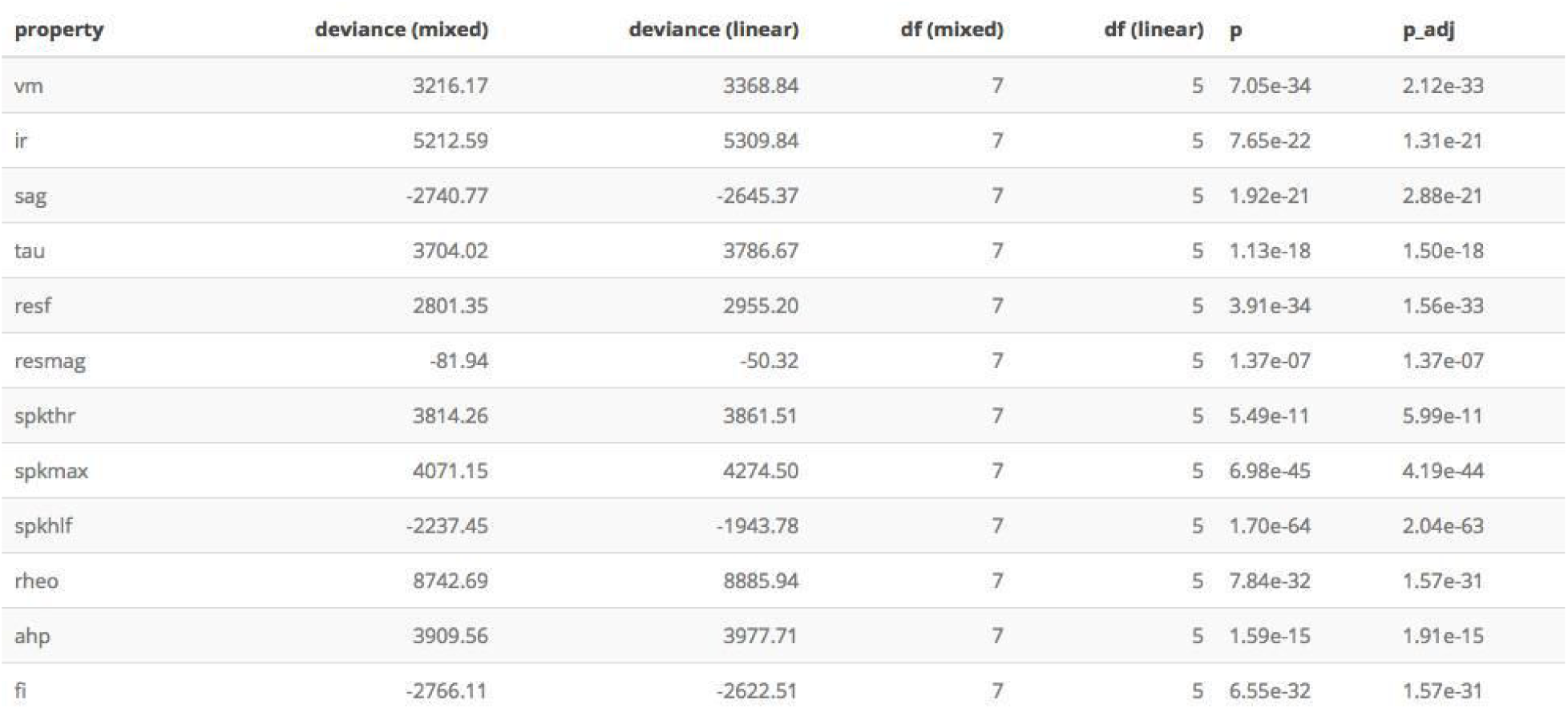
Inter-animal differences in electrophysiological features remain after accounting for housing. Results from comparison of mixed effect model incorporating dorsoventral location and housing, with an equivalent linear model. The significance estimate (p) is calculated using a chi squared test and adjusted for multiple comparisons (p_adj) using the Benjamini and Hochberg method.

**Supplemental Table 5.**
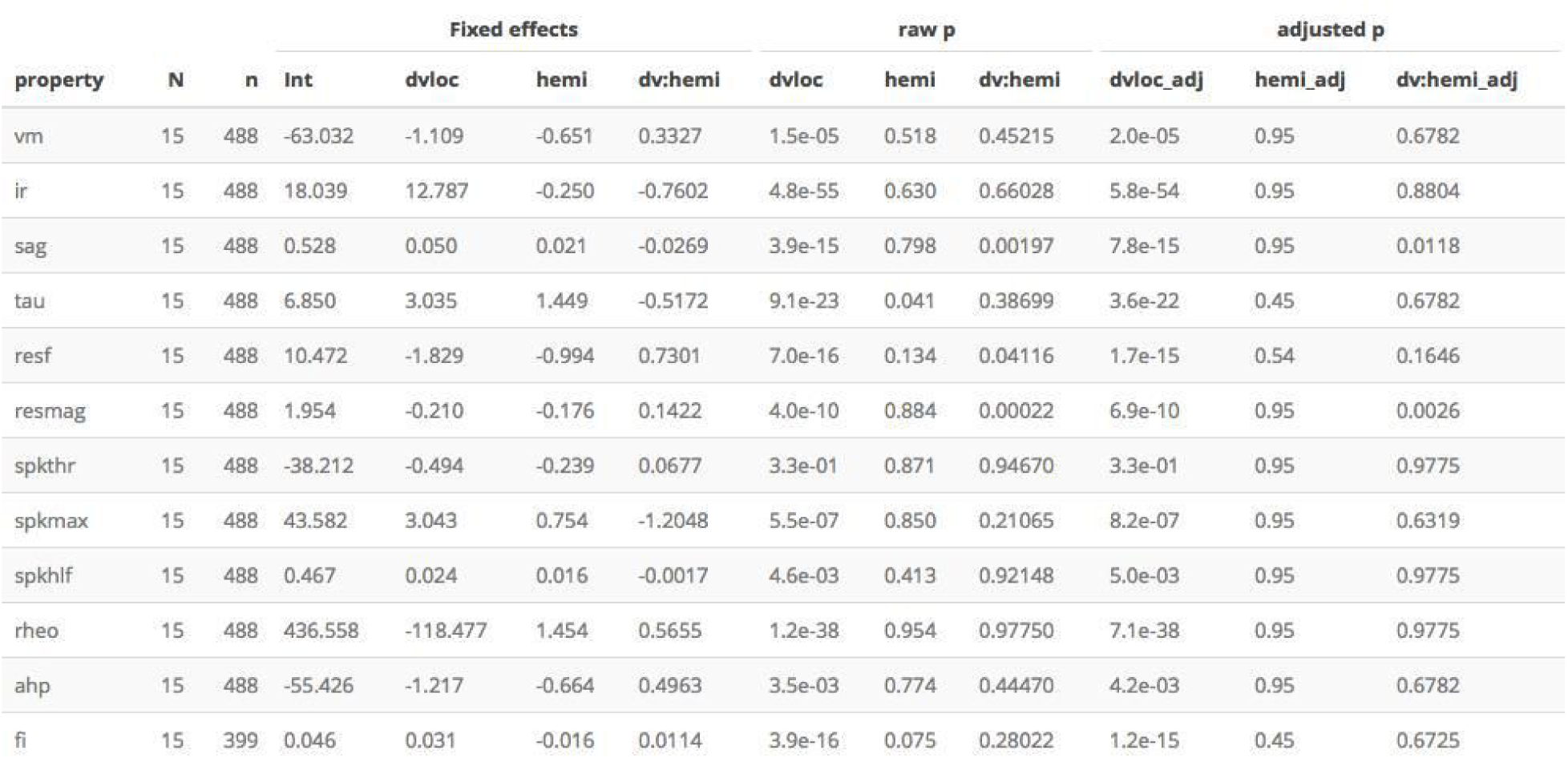
Dependence of SC properties on hemisphere. We did not find significant effects of brain hemisphere on any features except for the relationship between dorsoventral location and sag. Significance estimates for the effects of dorsoventral position (dvloc), brain hemisphere (hemi) and interactions between dorsoventral position and hemisphere (dvloc:hemi) were estimated using type II ANOVA and Wald *χ*^2^ test from fits to mixed models containing age and location as fixed effects and animal identity as random effects. Initial significance estimates (raw p) were adjusted for multiple comparisons (adjusted p) using the Benjamini and Hochberg method.

**Supplemental Table 6.**
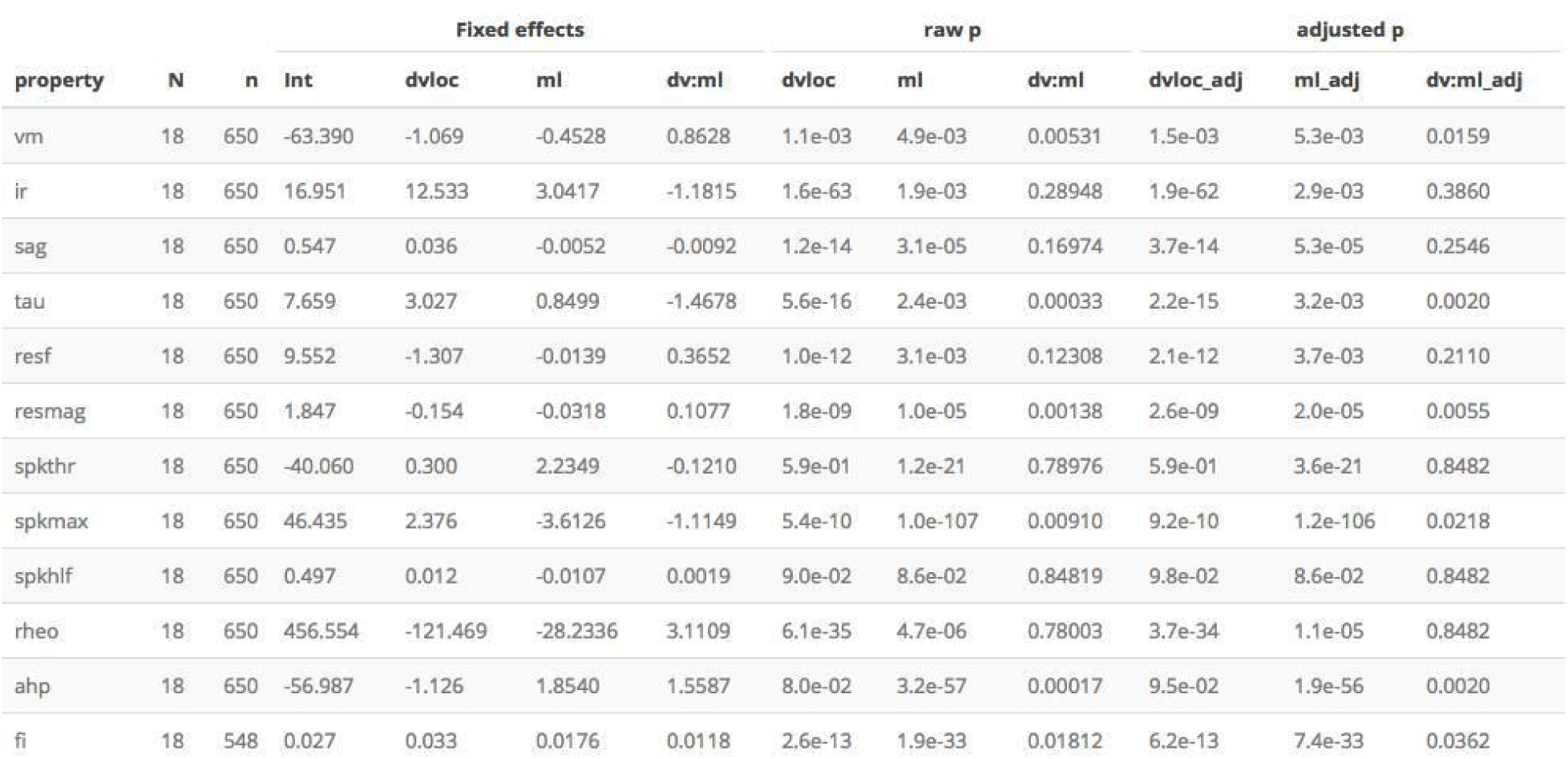
Dependence of SC properties on mediolateral position. Mediolateral as well as dorsoventral position has been reported to determine the sub-threshold electrophysiological features of SCs (Canto and Witter, 2012). We found significant effects of mediolateral position on all measured electrophysiological features. However, the sizes of the effects of mediolateral position on subthreshold features (vm, ir, sag, tau, resf, resmag, rheo) were much smaller than for dorsoventral position. In contrast, supra-threshold features (spkthr, spkmax, ahp) were more greatly affected by mediolateral position, with more medial neurons having a higher spike threshold, and lower amplitudes of the spike peak and after-hyperpolarization. Fixed effects are the intercept and slope coefficients for mixed models containing dorsoventral and mediolateral location as fixed effects and animal identity as random effects Significance estimates for the effects of dorsoventral position (dvloc), mediolateral position (ml) and interactions between dorsoventral position and mediolateral position (dvloc:ml) are estimated using type II ANOVA and Wald *χ*^2^ test from the fits of the mixed models. Initial significance estimates (raw p) were adjusted for multiple comparisons (adjusted p) using the Benjamini and Hochberg method.

**Supplemental Table 7.**
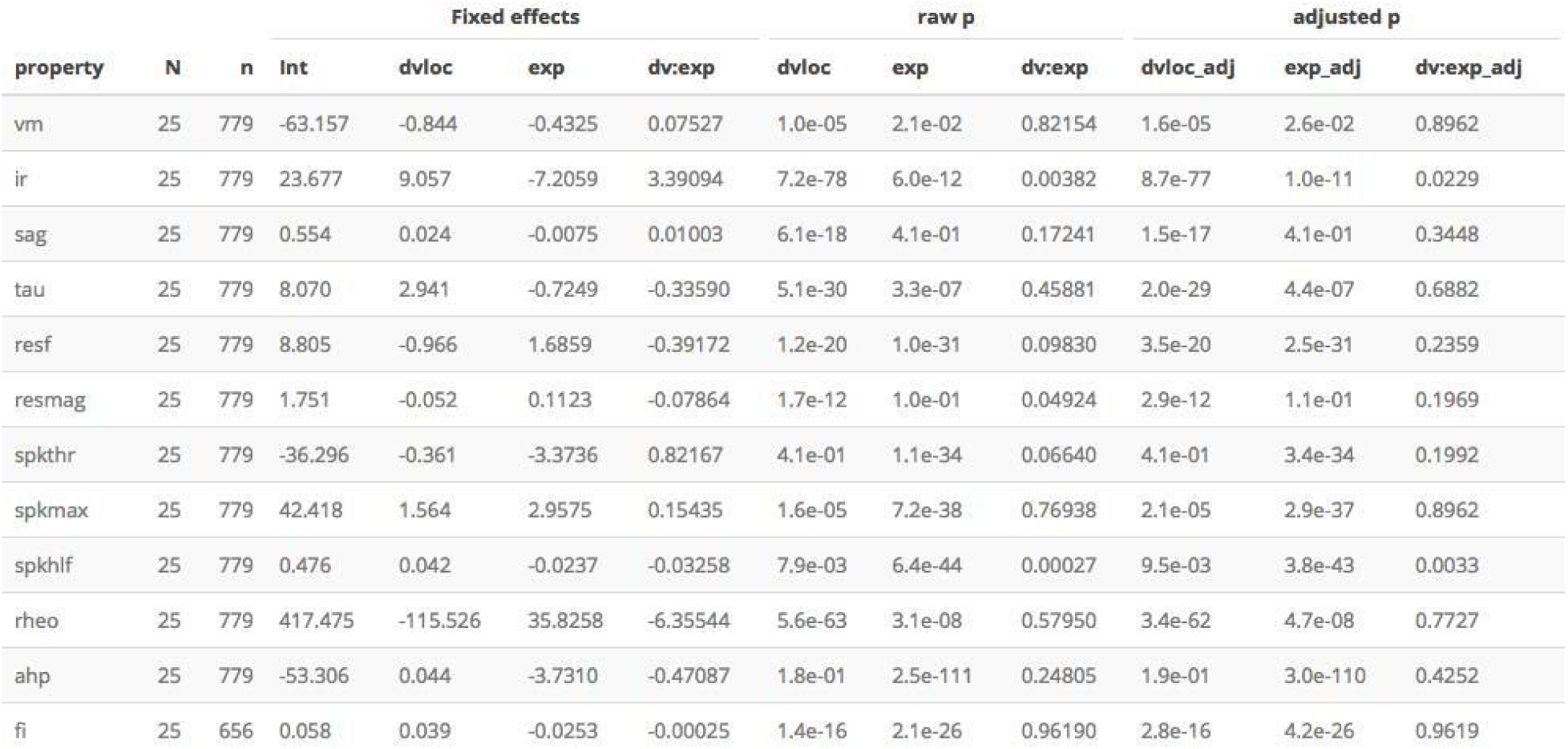
Dependence of SC properties on experimenter. We found that for many electrophysiological features the identity of the experimenter affected the intercept, but not the slope, of their relationship with dorsoventral position. All features except for spike threshold nevertheless followed a dorsoventral organisation after accounting for the experimenter. Significance estimates for the effects of dorsoventral position (dvloc), experimenter (exp) and interactions between dorsoventral position and experimenter (dvloc:exp) were estimated using type II ANOVA and Wald *χ*^2^ test from fits to mixed models containing age and location as fixed effects and animal identity as random effects. Initial significance estimates (raw p) were adjusted for multiple comparisons (adjusted p) using the Benjamini and Hochberg method.

**Supplemental Table 8.**
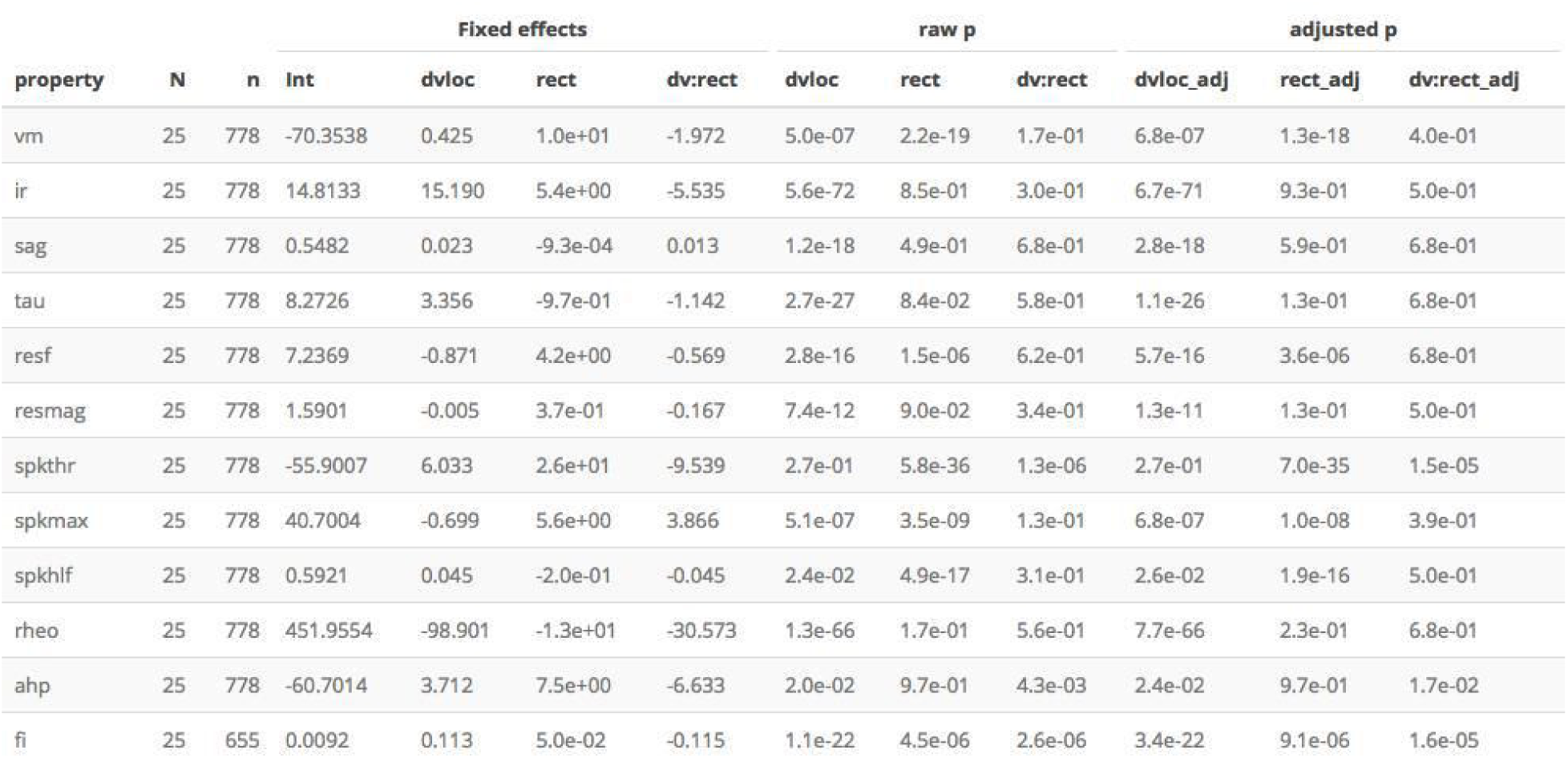
Dependence of SC properties on time since slice preparation. We anticipated that the interval between slice preparation and recording may influence measured electrophysiological features. Consistent with our expectation, analyses of the data were consistent with changes to some electrophysiological features of SCs with time since slice preparation, but dorsoventral gradients could not be explained by these changes (see also Supplemental Table 8 below). Significance estimates for the effects of dorsoventral position (dvloc), time since slice preparation (rect) and interactions between dorsoventral position and experimenter (dvloc:rect) estimated using type II ANOVA and Wald *χ*^2^ test from fits to mixed models containing age and location as fixed effects and animal identity as random effects. Initial significance estimates (raw p) were adjusted for multiple comparisons (adjusted p) using the Benjamini and Hochberg method.

| property | N | n | Int | Fixed effects |  |  | raw p |  |  | adjusted p |  |  |
| --- | --- | --- | --- | --- | --- | --- | --- | --- | --- | --- | --- | --- |
|  |  |  |  | dvloc | rect | dv:rect | dvloc | rect | dv:rect | dvloc_adj | rect_adj | dv:rect_adj |
| vm | 25 | 778 | -70.3538 | 0.425 | 1.0e+01 | -1.972 | 5.0e-07 | 2.2e-19 | 1.7e-01 | 6.8e-07 | 1.3e-18 | 4.0e-01 |
| ir | 25 | 778 | 14.8133 | 15.190 | 5.4e+00 | -5.535 | 5.6e-72 | 8.5e-01 | 3.0e-01 | 6.7e-71 | 9.3e-01 | 5.0e-01 |
| sag | 25 | 778 | 0.5482 | 0.023 | -9.3e-04 | 0.013 | 1.2e-18 | 4.9e-01 | 6.8e-01 | 2.8e-18 | 5.9e-01 | 6.8e-01 |
| tau | 25 | 778 | 8.2726 | 3.356 | -9.7e-01 | -1.142 | 2.7e-27 | 8.4e-02 | 5.8e-01 | 1.1e-26 | 1.3e-01 | 6.8e-01 |
| resf | 25 | 778 | 7.2369 | -0.871 | 4.2e+00 | -0.569 | 2.8e-16 | 1.5e-06 | 6.2e-01 | 5.7e-16 | 3.6e-06 | 6.8e-01 |
| resmag | 25 | 778 | 1.5901 | -0.005 | 3.7e-01 | -0.167 | 7.4e-12 | 9.0e-02 | 3.4e-01 | 1.3e-11 | 1.3e-01 | 5.0e-01 |
| spkthr | 25 | 778 | -55.9007 | 6.033 | 2.6e+01 | -9.539 | 2.7e-01 | 5.8e-36 | 1.3e-06 | 2.7e-01 | 7.0e-35 | 1.5e-05 |
| spkmax | 25 | 778 | 40.7004 | -0.699 | 5.6e+00 | 3.866 | 5.1e-07 | 3.5e-09 | 1.3e-01 | 6.8e-07 | 1.0e-08 | 3.9e-01 |
| spkhlf | 25 | 778 | 0.5921 | 0.045 | -2.0e-01 | -0.045 | 2.4e-02 | 4.9e-17 | 3.1e-01 | 2.6e-02 | 1.9e-16 | 5.0e-01 |
| rheo | 25 | 778 | 451.9554 | -98.901 | -1.3e+01 | -30.573 | 1.3e-66 | 1.7e-01 | 5.6e-01 | 7.7e-66 | 2.3e-01 | 6.8e-01 |
| ahp | 25 | 778 | -60.7014 | 3.712 | 7.5e+00 | -6.633 | 2.0e-02 | 9.7e-01 | 4.3e-03 | 2.4e-02 | 9.7e-01 | 1.7e-02 |
| fi | 25 | 655 | 0.0092 | 0.113 | 5.0e-02 | -0.115 | 1.1e-22 | 4.5e-06 | 2.6e-06 | 3.4e-22 | 9.1e-06 | 1.6e-05 |

**Supplemental Table 9.** Dependence of SC properties on direction in which sequential recordings are made. In anticipation of effects of the time since slice preparation on electrophysiological features of SCS, we varied between experimenters and experimental days the direction along the dorsoventral axis from which consecutive recordings were made (see Methods). Consistent with effects of time on electrophysiological features (see Supplemental Table 7 above), we found that the direction in which sequential recordings were made influence the slope, but not the intercept of several electrophysiological features. Significance estimates for the effects of dorsoventral position (dvloc), direction in which sequential recordings were made (dir) and interactions between dorsoventral position and recording direction (dvloc:dir) estimated using type II ANOVA and Wald *χ*^2^ test from fits to mixed models containing age and location as fixed effects and animal identity as random effects. Initial significance estimates (raw p) were adjusted for multiple comparisons (adjusted p) using the Benjamini and Hochberg method.

| property | N | n | Fixed effects |  |  |  | raw p |  |  | adjusted p |  |  |
| --- | --- | --- | --- | --- | --- | --- | --- | --- | --- | --- | --- | --- |
|  |  |  | Int | dvloc | dir | dv:dir | dvloc | dir | dv:dir | dvloc_adj | dir_adj | dv:dir_adj |
| vm | 18 | 650 | -64.127 | -0.063 | 1.351 | -1.3535 | 2.8e-04 | 0.509 | 2.4e-05 | 3.7e-04 | 0.61 | 9.4e-05 |
| ir | 18 | 650 | 18.337 | 11.758 | 0.427 | 0.1689 | 8.7e-64 | 0.374 | 8.8e-01 | 1.0e-62 | 0.57 | 9.2e-01 |
| sag | 18 | 650 | 0.534 | 0.040 | 0.021 | -0.0164 | 1.3e-16 | 0.411 | 1.9e-02 | 3.1e-16 | 0.57 | 3.8e-02 |
| tau | 18 | 650 | 7.975 | 2.269 | 0.091 | 0.2595 | 2.5e-20 | 0.144 | 5.4e-01 | 1.0e-19 | 0.49 | 7.2e-01 |
| resf | 18 | 650 | 9.372 | -0.905 | 0.443 | -0.5214 | 1.9e-12 | 0.429 | 3.7e-02 | 3.8e-12 | 0.57 | 6.3e-02 |
| resmag | 18 | 650 | 1.846 | -0.104 | -0.030 | -0.0035 | 1.0e-08 | 0.122 | 9.2e-01 | 1.7e-08 | 0.49 | 9.2e-01 |
| spkthr | 18 | 650 | -40.893 | 1.947 | 4.997 | -4.2461 | 5.2e-01 | 0.219 | 1.0e-23 | 5.2e-01 | 0.49 | 1.2e-22 |
| spkmax | 18 | 650 | 43.529 | 3.044 | 2.134 | -1.7908 | 5.0e-08 | 0.556 | 2.4e-03 | 7.4e-08 | 0.61 | 5.9e-03 |
| spkhlf | 18 | 650 | 0.511 | -0.012 | -0.042 | 0.0491 | 1.2e-01 | 0.059 | 2.8e-06 | 1.3e-01 | 0.49 | 1.7e-05 |
| rheo | 18 | 650 | 439.066 | -119.612 | 10.708 | -1.5603 | 2.0e-43 | 0.168 | 9.0e-01 | 1.2e-42 | 0.49 | 9.2e-01 |
| ahp | 18 | 650 | -56.018 | -0.474 | 0.408 | -0.4158 | 1.1e-02 | 0.872 | 4.0e-01 | 1.3e-02 | 0.87 | 6.0e-01 |
| fi | 18 | 548 | 0.047 | 0.028 | -0.023 | 0.0189 | 7.5e-18 | 0.245 | 6.5e-04 | 2.2e-17 | 0.49 | 1.9e-03 |

**Supplemental Table 10.** Interanimal differences in extended models. Results from comparison of mixed effect model incorporating dorsoventral location, housing, mediolateral position, experimenter identity and direction in which recordings were obtained with an equivalent linear model. Data are from animals between 32 and 45 days old. The significance estimate (p) is calculated using a chi squared test and adjusted for multiple comparisons (p_adj) using the Benjamini and Hochberg method.

| property | deviance (mixed) | deviance (linear) | df (mixed) | df (linear) | p | p_adj |
| --- | --- | --- | --- | --- | --- | --- |
| vm | 2636.81 | 2789.34 | 13 | 11 | 7.56e-34 | 3.02e-33 |
| ir | 4315.73 | 4407.44 | 13 | 11 | 1.22e-20 | 1.62e-20 |
| sag | -2293.74 | -2212.49 | 13 | 11 | 2.27e-18 | 2.72e-18 |
| tau | 2997.16 | 3122.25 | 13 | 11 | 6.86e-28 | 1.37e-27 |
| resf | 2122.53 | 2266.61 | 13 | 11 | 5.16e-32 | 1.55e-31 |
| resmag | -115.70 | -80.54 | 13 | 11 | 2.31e-08 | 2.31e-08 |
| spkthr | 2936.17 | 2974.72 | 13 | 11 | 4.25e-09 | 4.63e-09 |
| spkmax | 3008.01 | 3239.16 | 13 | 11 | 6.40e-51 | 7.67e-50 |
| spkhlf | -2067.19 | -1911.08 | 13 | 11 | 1.26e-34 | 7.55e-34 |
| rheo | 7247.31 | 7388.78 | 13 | 11 | 1.91e-31 | 4.58e-31 |
| ahp | 2832.78 | 2943.86 | 13 | 11 | 7.58e-25 | 1.14e-24 |
| fi | -2473.53 | -2349.35 | 13 | 11 | 1.08e-27 | 1.85e-27 |

**Supplemental Table 11.**
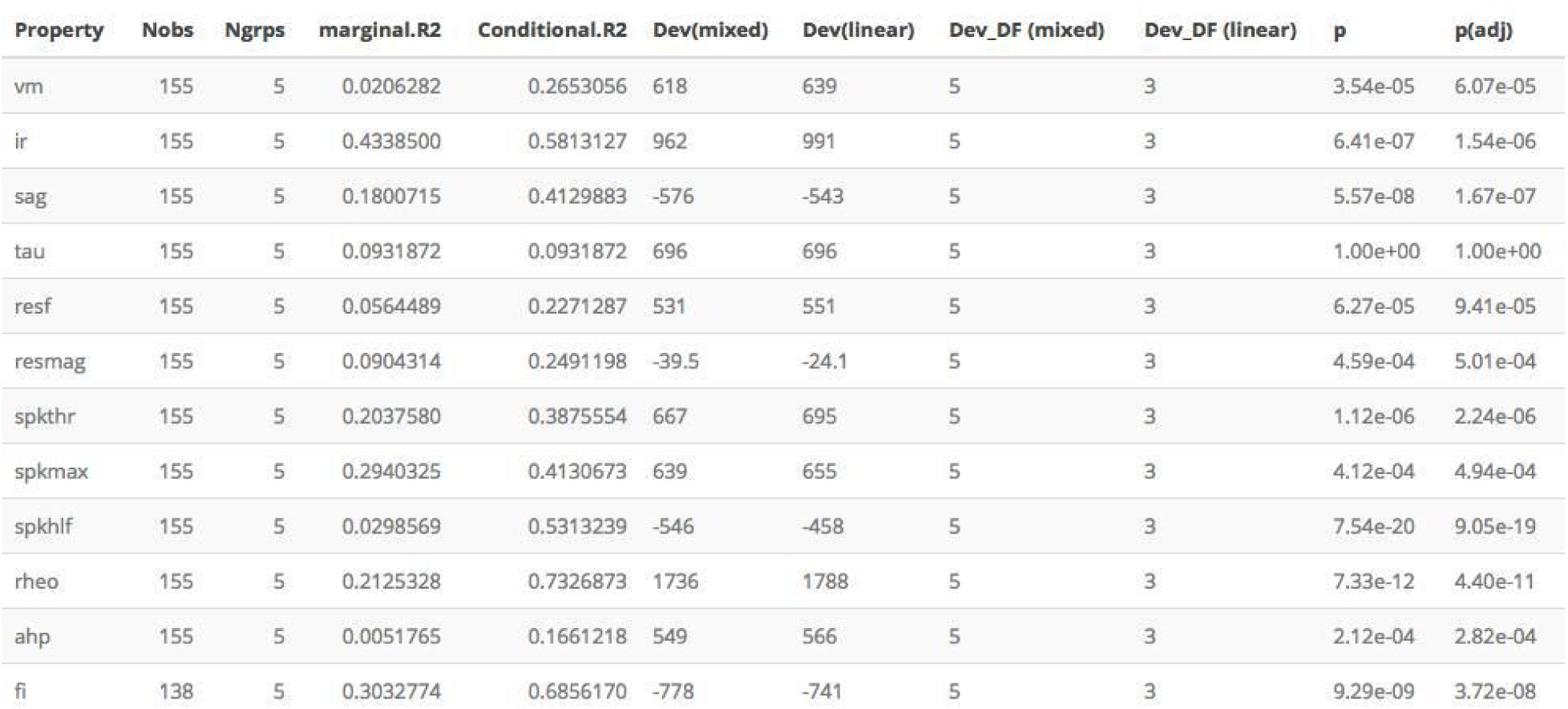
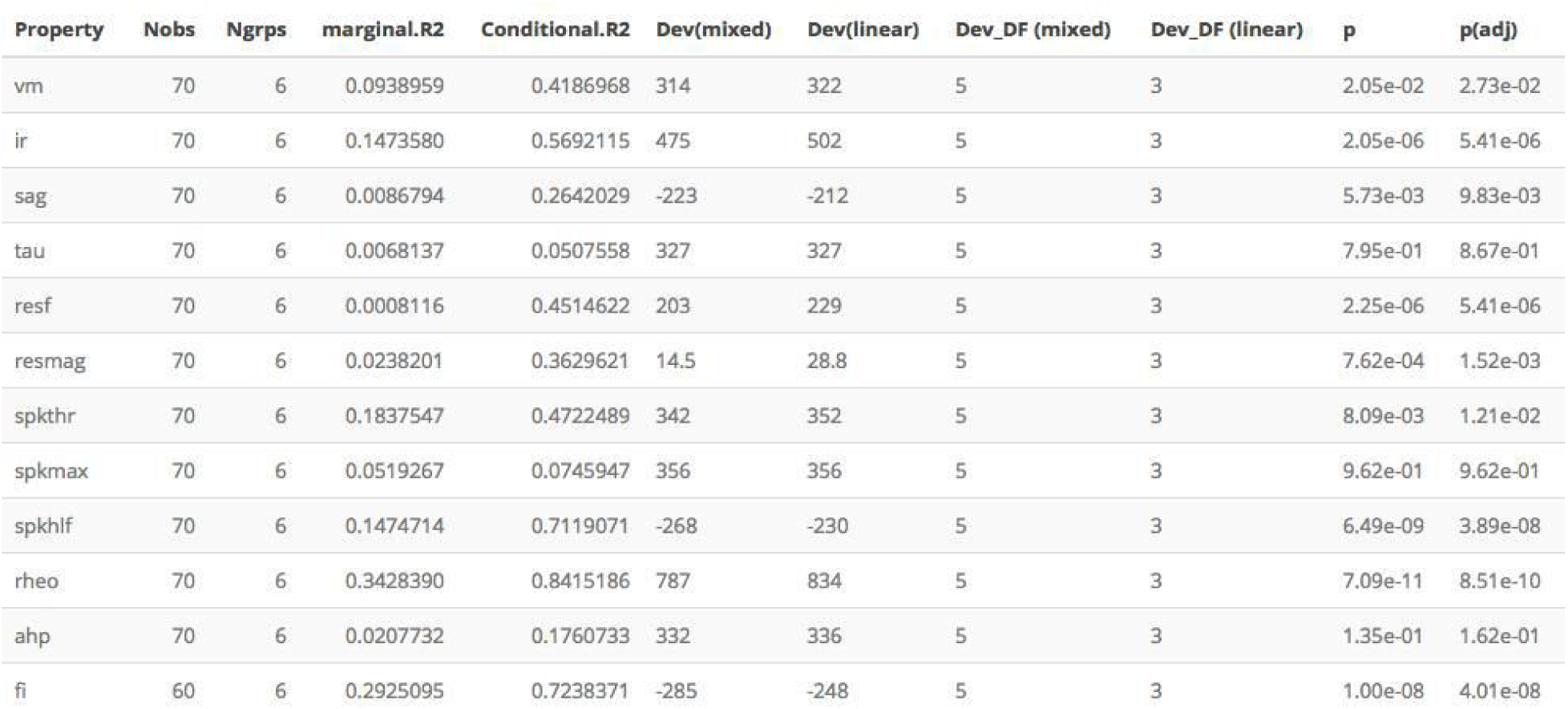
Interanimal differences in models fit to minimal datasets. Results from comparison of mixed effect models with dorsoventral location as a fixed effect and animal identity as a random effect using a minimal datasets obtained by either HP (upper) or DG (lower). Data are from animals between 32 and 45 days old. Because of the smaller size of these datasets the statistical power to detect inter-animal variation is reduced. Nevertheless, in these analyses the conditional R^2^ of the mixed model fit was again substantially higher than the marginal R^2^, and most (9/12) features were better fit by a mixed model compared to a corresponding linear model in both datasets.

| Property | Nobs | Ngrps | marginal.R2 | Conditional.R2 | Dev(mixed) | Dev(linear) | Dev_DF (mixed) | Dev_DF (linear) | p | p(adj) |
| --- | --- | --- | --- | --- | --- | --- | --- | --- | --- | --- |
| vm | 155 | 5 | 0.0206282 | 0.2653056 | 618 | 639 | 5 | 3 | 3.54e-05 | 6.07e-05 |
| ir | 155 | 5 | 0.4338500 | 0.5813127 | 962 | 991 | 5 | 3 | 6.41e-07 | 1.54e-06 |
| sag | 155 | 5 | 0.1800715 | 0.4129883 | -576 | -543 | 5 | 3 | 5.57e-08 | 1.67e-07 |
| tau | 155 | 5 | 0.0931872 | 0.0931872 | 696 | 696 | 5 | 3 | 1.00e+00 | 1.00e+00 |
| resf | 155 | 5 | 0.0564489 | 0.2271287 | 531 | 551 | 5 | 3 | 6.27e-05 | 9.41e-05 |
| resmag | 155 | 5 | 0.0904314 | 0.2491198 | -39.5 | -24.1 | 5 | 3 | 4.59e-04 | 5.01e-04 |
| spkthr | 155 | 5 | 0.2037580 | 0.3875554 | 667 | 695 | 5 | 3 | 1.12e-06 | 2.24e-06 |
| spkmax | 155 | 5 | 0.2940325 | 0.4130673 | 639 | 655 | 5 | 3 | 4.12e-04 | 4.94e-04 |
| spkhlf | 155 | 5 | 0.0298569 | 0.5313239 | -546 | -458 | 5 | 3 | 7.54e-20 | 9.05e-19 |
| rheo | 155 | 5 | 0.2125328 | 0.7326873 | 1736 | 1788 | 5 | 3 | 7.33e-12 | 4.40e-11 |
| ahp | 155 | 5 | 0.0051765 | 0.1661218 | 549 | 566 | 5 | 3 | 2.12e-04 | 2.82e-04 |
| fi | 138 | 5 | 0.3032774 | 0.6856170 | -778 | -741 | 5 | 3 | 9.29e-09 | 3.72e-08 |
| vm | 70 | 6 | 0.0938959 | 0.4186968 | 314 | 322 | 5 | 3 | 2.05e-02 | 2.73e-02 |
| ir | 70 | 6 | 0.1473580 | 0.5692115 | 475 | 502 | 5 | 3 | 2.05e-06 | 5.41e-06 |
| sag | 70 | 6 | 0.0086794 | 0.2642029 | -223 | -212 | 5 | 3 | 5.73e-03 | 9.83e-03 |
| tau | 70 | 6 | 0.0068137 | 0.0507558 | 327 | 327 | 5 | 3 | 7.95e-01 | 8.67e-01 |
| resf | 70 | 6 | 0.0008116 | 0.4514622 | 203 | 229 | 5 | 3 | 2.25e-06 | 5.41e-06 |
| resmag | 70 | 6 | 0.0238201 | 0.3629621 | 14.5 | 28.8 | 5 | 3 | 7.62e-04 | 1.52e-03 |
| spkthr | 70 | 6 | 0.1837547 | 0.4722489 | 342 | 352 | 5 | 3 | 8.09e-03 | 1.21e-02 |
| spkmax | 70 | 6 | 0.0519267 | 0.0745947 | 356 | 356 | 5 | 3 | 9.62e-01 | 9.62e-01 |
| spkhlf | 70 | 6 | 0.1474714 | 0.7119071 | -268 | -230 | 5 | 3 | 6.49e-09 | 3.89e-08 |
| rheo | 70 | 6 | 0.3428390 | 0.8415186 | 787 | 834 | 5 | 3 | 7.09e-11 | 8.51e-10 |
| ahp | 70 | 6 | 0.0207732 | 0.1760733 | 332 | 336 | 5 | 3 | 1.35e-01 | 1.62e-01 |
| fi | 60 | 6 | 0.2925095 | 0.7238371 | -285 | -248 | 5 | 3 | 1.00e-08 | 4.01e-08 |

**Supplemental Table 12.** Electrophysiological features and the number of recorded neurons. We considered whether variation in tissue quality as a contributor to inter-animal variability. To minimise this possibility we standardised our procedures for tissue preparation (see Methods), such that slices were of consistent high quality as assessed by low numbers of unhealthy cells and by visualisation of soma and dendrites of neurons in the slice. We further reasoned that if the condition of the slices differed between animals, then in better quality slices it would be easier to record from more neurons, in which case any features that depend on tissue quality would correlate with the number of recorded neurons. We found that the majority (10/12) of electrophysiological features were not significantly (p > 0.2) associated with the number of recorded neurons. Significance estimates for the effects of dorsoventral position (dvloc), number of recorded neurons (counts) and interactions between dorsoventral position and number of recorded neurons (dvloc:counts) estimated using type II ANOVA and Wald *χ*^2^ test from fits to mixed models containing age and location as fixed effects and animal identity as random effects. Initial significance estimates (raw p) were adjusted for multiple comparisons (adjusted p) using the Benjamini and Hochberg method.

| property | N | n | Fixed effects |  |  |  | raw p |  |  | adjusted p |  |  |
| --- | --- | --- | --- | --- | --- | --- | --- | --- | --- | --- | --- | --- |
|  |  |  | Int | dvloc | counts | dv:counts | dvloc | counts | dv:counts | dvloc_adj | counts_adj | dv:counts_adj |
| vm | 25 | 779 | -63.262 | -1.334 | -0.00470 | 1.5e-02 | 7.2e-06 | 0.72309 | 0.384 | 9.7e-06 | 0.9641 | 0.58 |
| ir | 25 | 779 | 20.165 | 8.240 | -0.04768 | 9.6e-02 | 1.2e-75 | 0.61560 | 0.113 | 1.4e-74 | 0.9234 | 0.23 |
| sag | 25 | 779 | 0.568 | 0.010 | -0.00055 | 6.2e-04 | 8.3e-20 | 0.98892 | 0.080 | 2.5e-19 | 0.9889 | 0.23 |
| tau | 25 | 779 | 5.964 | 3.987 | 0.04659 | -3.9e-02 | 8.6e-28 | 0.34149 | 0.092 | 3.4e-27 | 0.9234 | 0.23 |
| resf | 25 | 779 | 12.067 | -1.880 | -0.06141 | 1.8e-02 | 1.2e-18 | 0.00231 | 0.194 | 2.8e-18 | 0.0139 | 0.33 |
| resmag | 25 | 779 | 1.777 | -0.066 | 0.00166 | -1.3e-03 | 5.3e-12 | 0.83262 | 0.433 | 9.1e-12 | 0.9889 | 0.58 |
| spkthr | 25 | 779 | -38.810 | 0.278 | 0.00217 | -3.3e-03 | 6.7e-01 | 0.98673 | 0.925 | 6.7e-01 | 0.9889 | 0.97 |
| spkmax | 25 | 779 | 44.731 | 0.051 | -0.00543 | 5.4e-02 | 1.5e-07 | 0.46922 | 0.117 | 2.2e-07 | 0.9234 | 0.23 |
| spkhlf | 25 | 779 | 0.327 | 0.064 | 0.00408 | -1.4e-03 | 1.4e-02 | 0.00022 | 0.019 | 1.6e-02 | 0.0027 | 0.22 |
| rheo | 25 | 779 | 388.527 | -73.875 | 1.56603 | -1.3e+00 | 8.3e-74 | 0.45667 | 0.043 | 5.0e-73 | 0.9234 | 0.23 |
| ahp | 25 | 779 | -56.673 | -0.185 | 0.02539 | -1.2e-02 | 2.7e-02 | 0.59590 | 0.659 | 2.9e-02 | 0.9234 | 0.79 |
| fi | 25 | 656 | 0.052 | 0.036 | -0.00032 | 1.3e-05 | 5.5e-18 | 0.32090 | 0.973 | 1.1e-17 | 0.9234 | 0.97 |

**Supplemental Table 13.**
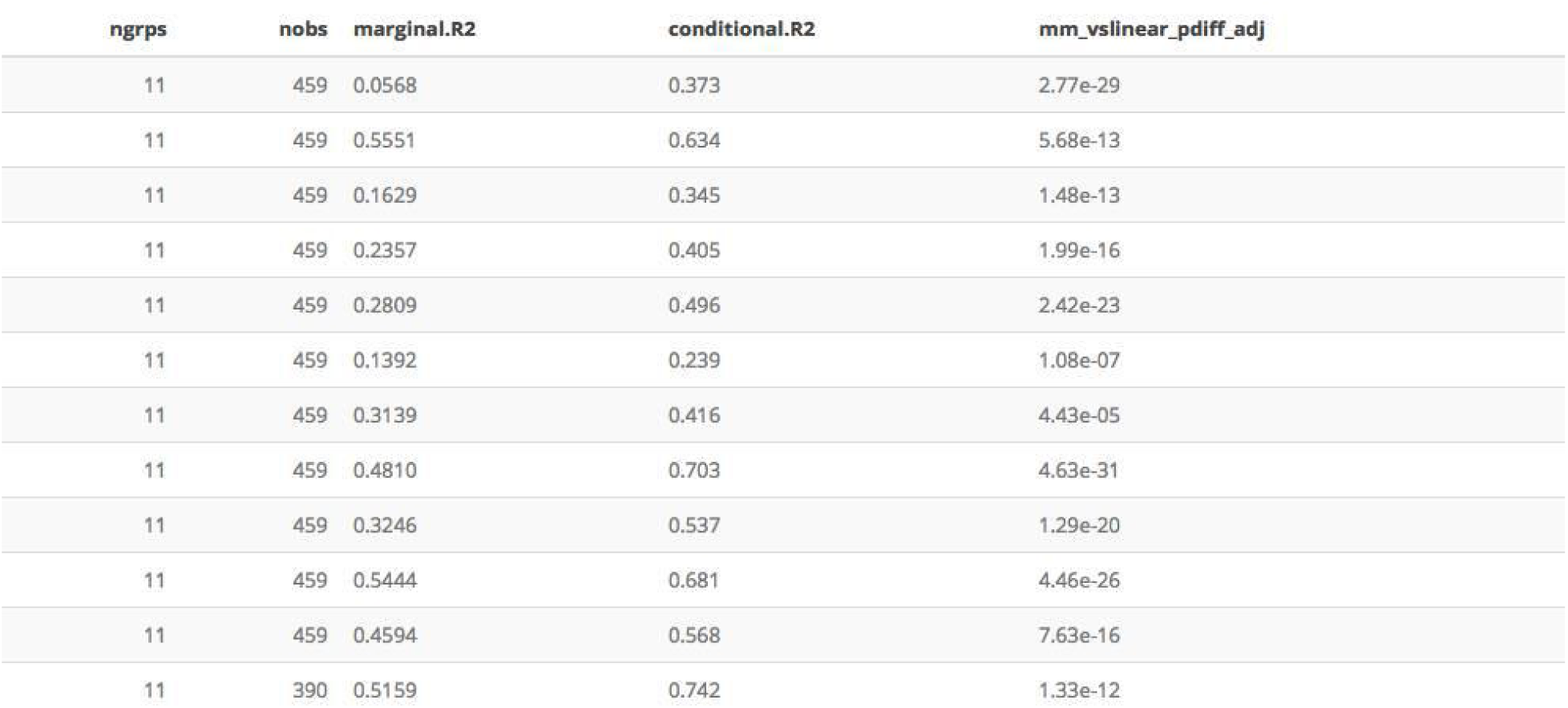
Interanimal differences for experiments with > 35 recorded neurons. To further assess the robustness of our results to possible variations in slice quality, we repeated analyses of inter-animal differences but focussing only on data from animals for which > 35 recordings were made (N = 11, n = 459). Comparison of marginal and conditional R^2^ values continued to indicate substantial inter-animal variance, and fits obtained with mixed models remained significantly different to fits that did not account for animal identity (p < 4.4 × 10^-5^). Analyses are as for Supplemental Table 1, but are restricted to experiments in which > 35 neurons were recorded from.

**Supplemental Table 14.** Dependence of principal components on dorsoventral position and animal identity. Analyses are as described for Table 1, but are applied to principal components of the electrophysiological features of SCs.

| Feature | Slope | p (slope) | Marginal R2 | Conditional R2 | Slope (min) | Slope (max) | p (vs linear) |
| --- | --- | --- | --- | --- | --- | --- | --- |
| PC1 | -2.43035 | 1.09e-15 | 0.49545 | 0.73250 | -3.16510 | -2.07369 | 3.86e-37 |
| PC2 | 0.95272 | 1.05e-04 | 0.09299 | 0.80751 | -1.07716 | 2.11870 | 2.20e-97 |
| PC3 | 0.25632 | 1.73e-01 | 0.01187 | 0.51214 | -0.43529 | 1.11057 | 2.96e-38 |
| PC4 | -0.26627 | 2.53e-01 | 0.00941 | 0.79049 | -1.47230 | 1.06857 | 4.45e-63 |
| PC5 | -0.11523 | 4.01e-01 | 0.00390 | 0.43605 | -0.49011 | 0.44712 | 3.10e-24 |
| PC6 | -0.01593 | 9.03e-01 | 0.00006 | 0.64068 | -1.12576 | 1.08528 | 7.98e-31 |
| PC7 | -0.06545 | 4.90e-01 | 0.00250 | 0.05729 | -0.18349 | 0.07602 | 2.94e-02 |
| PC8 | -0.01013 | 9.03e-01 | 0.00007 | 0.23751 | -0.31346 | 0.24927 | 3.49e-10 |
| PC9 | -0.13547 | 3.95e-02 | 0.01617 | 0.18557 | -0.22543 | 0.08319 | 4.98e-12 |
| PC10 | -0.01408 | 9.03e-01 | 0.00031 | 0.04810 | -0.02747 | -0.00033 | 8.19e-02 |
| PC11 | -0.00545 | 9.03e-01 | 0.00007 | 0.02613 | -0.00794 | -0.00383 | 2.74e-01 |
| PC12 | -0.07795 | 1.50e-02 | 0.02402 | 0.08892 | -0.19333 | 0.01590 | 7.02e-04 |

**Supplemental Table 15.** Dependence of principal components of SC properties on housing. Analyses are as described for Supplemental Table 3, but are applied to principal components of the electrophysiological features of SCs.

| property | N | n | Fixed effects |  |  |  | raw p |  |  | adjusted p |  |  |
| --- | --- | --- | --- | --- | --- | --- | --- | --- | --- | --- | --- | --- |
|  |  |  | Int | dvloc | housing | dv:housing | dvloc | housing | dv:housing | dvloc_adj | housing_adj | dv:housing_adj |
| PC1 | 25 | 487 | 2.055 | -2.373 | 0.9582 | -0.0754 | 2.7e-86 | 0.0098 | 0.77164 | 3.3e-85 | 0.039 | 0.8890 |
| PC2 | 25 | 487 | -1.859 | 1.067 | 1.2929 | -0.1590 | 2.1e-07 | 0.0045 | 0.68640 | 1.2e-06 | 0.027 | 0.8890 |
| PC3 | 25 | 487 | -0.157 | 0.301 | -0.2277 | -0.0681 | 6.3e-02 | 0.4571 | 0.81489 | 1.5e-01 | 0.609 | 0.8890 |
| PC4 | 25 | 487 | 0.754 | -0.369 | -0.6951 | 0.1501 | 1.1e-01 | 0.1441 | 0.67220 | 2.2e-01 | 0.247 | 0.8890 |
| PC5 | 25 | 487 | 0.284 | -0.114 | -0.1393 | -0.0016 | 2.3e-01 | 0.6192 | 0.99353 | 4.0e-01 | 0.681 | 0.9935 |
| PC6 | 25 | 487 | 0.436 | -0.238 | -0.4509 | 0.3286 | 8.8e-01 | 0.2440 | 0.21869 | 9.4e-01 | 0.366 | 0.6561 |
| PC7 | 25 | 487 | 0.398 | -0.279 | -0.4526 | 0.3226 | 3.1e-01 | 0.0991 | 0.02447 | 4.6e-01 | 0.238 | 0.0979 |
| PC8 | 25 | 487 | 0.277 | -0.057 | -0.3795 | 0.0706 | 8.7e-01 | 0.0037 | 0.59369 | 9.4e-01 | 0.027 | 0.8890 |
| PC9 | 25 | 487 | 0.134 | -0.177 | -0.0042 | 0.0658 | 1.3e-02 | 0.6239 | 0.56105 | 3.8e-02 | 0.681 | 0.8890 |
| PC10 | 25 | 487 | 0.097 | -0.138 | -0.1354 | 0.2118 | 9.4e-01 | 0.0430 | 0.00576 | 9.4e-01 | 0.129 | 0.0345 |
| PC11 | 25 | 487 | 0.045 | -0.039 | -0.0537 | 0.0535 | 8.8e-01 | 0.8677 | 0.41405 | 9.4e-01 | 0.868 | 0.8890 |
| PC12 | 25 | 487 | -0.076 | 0.046 | 0.2241 | -0.1870 | 3.7e-03 | 0.1242 | 0.00056 | 1.5e-02 | 0.247 | 0.0067 |

**Supplemental Table 16.** Dependence of principal components on animal identity in models that account for housing. Analyses are as for Supplemental Table 10, but are applied to principal components of the electrophysiological features of SCs.

| component | deviance (mixed) | deviance (linear) | df (mixed) | df (linear) | p | p_adj |
| --- | --- | --- | --- | --- | --- | --- |
| PC1 | 1463.86 | 1596.78 | 7 | 5 | 1.37e-29 | 3.28e-29 |
| PC2 | 1260.20 | 1658.40 | 7 | 5 | 3.40e-87 | 4.08e-86 |
| PC3 | 1394.81 | 1567.87 | 7 | 5 | 2.63e-38 | 1.05e-37 |
| PC4 | 1180.62 | 1441.66 | 7 | 5 | 2.07e-57 | 1.24e-56 |
| PC5 | 1217.87 | 1319.67 | 7 | 5 | 7.83e-23 | 1.57e-22 |
| PC6 | 1104.30 | 1237.28 | 7 | 5 | 1.33e-29 | 3.28e-29 |
| PC7 | 1070.53 | 1078.31 | 7 | 5 | 2.05e-02 | 2.46e-02 |
| PC8 | 904.79 | 929.29 | 7 | 5 | 4.78e-06 | 7.17e-06 |
| PC9 | 827.72 | 881.34 | 7 | 5 | 2.27e-12 | 3.89e-12 |
| PC10 | 590.98 | 592.69 | 7 | 5 | 4.25e-01 | 4.25e-01 |
| PC11 | 432.20 | 434.54 | 7 | 5 | 3.12e-01 | 3.40e-01 |
| PC12 | 116.24 | 128.36 | 7 | 5 | 2.34e-03 | 3.12e-03 |

